# Identification of the main barriers to Ku accumulation in chromatin

**DOI:** 10.1101/2024.01.03.574002

**Authors:** Madeleine Bossaert, Andrew Moreno, Antonio Peixoto, Marie-Jeanne Pillaire, Pauline Chanut, Philippe Frit, Patrick Calsou, Joseph John Loparo, Sébastien Britton

## Abstract

Repair of DNA double strand breaks by the non-homologous end-joining pathway is initiated by the binding of Ku to DNA ends. Given its high affinity for ends, multiple Ku proteins load onto linear DNAs in vitro. However, in cells, Ku loading is limited to ∼1-2 molecules per DNA end. The mechanisms enforcing this limit are currently unknown. Here we show that the catalytic subunit of the DNA-dependent protein kinase (DNA-PKcs), but not its protein kinase activity, is required to prevent excessive Ku entry into chromatin. Ku accumulation is further restricted by two mechanisms: a neddylation/FBXL12-dependent process which actively removes loaded Ku molecules throughout the cell cycle and a CtIP/ATM-dependent mechanism which operates in S-phase. Finally, we demonstrate that the misregulation of Ku loading leads to impaired transcription in the vicinity of DNA ends. Together our data shed light on the multiple layers of coordinated mechanisms operating to prevent Ku from invading chromatin and interfering with other DNA transactions.

**Highlights:**

- DNA-PKcs structurally blocks Ku sliding into chromatin in human & *Xenopus*
- A neddylation/FBXL12-dependent mechanism limits Ku accumulation on chromatin
- In S-phase, ATM/CtIP overcomes Ku accumulation
- In absence of DNA-PKcs, transcription at the DNA end vicinity is inhibited

**eTOC blurb:** The DNA end binding protein Ku can slide onto naked DNA but this is limited in cells. Using human cells and *Xenopus* egg extracts, DNA-PKcs is identified as the main structural barrier to Ku entry into chromatin, along with two active mechanisms which limit Ku accumulation in absence of DNA-PKcs.

**Graphical abstract:** 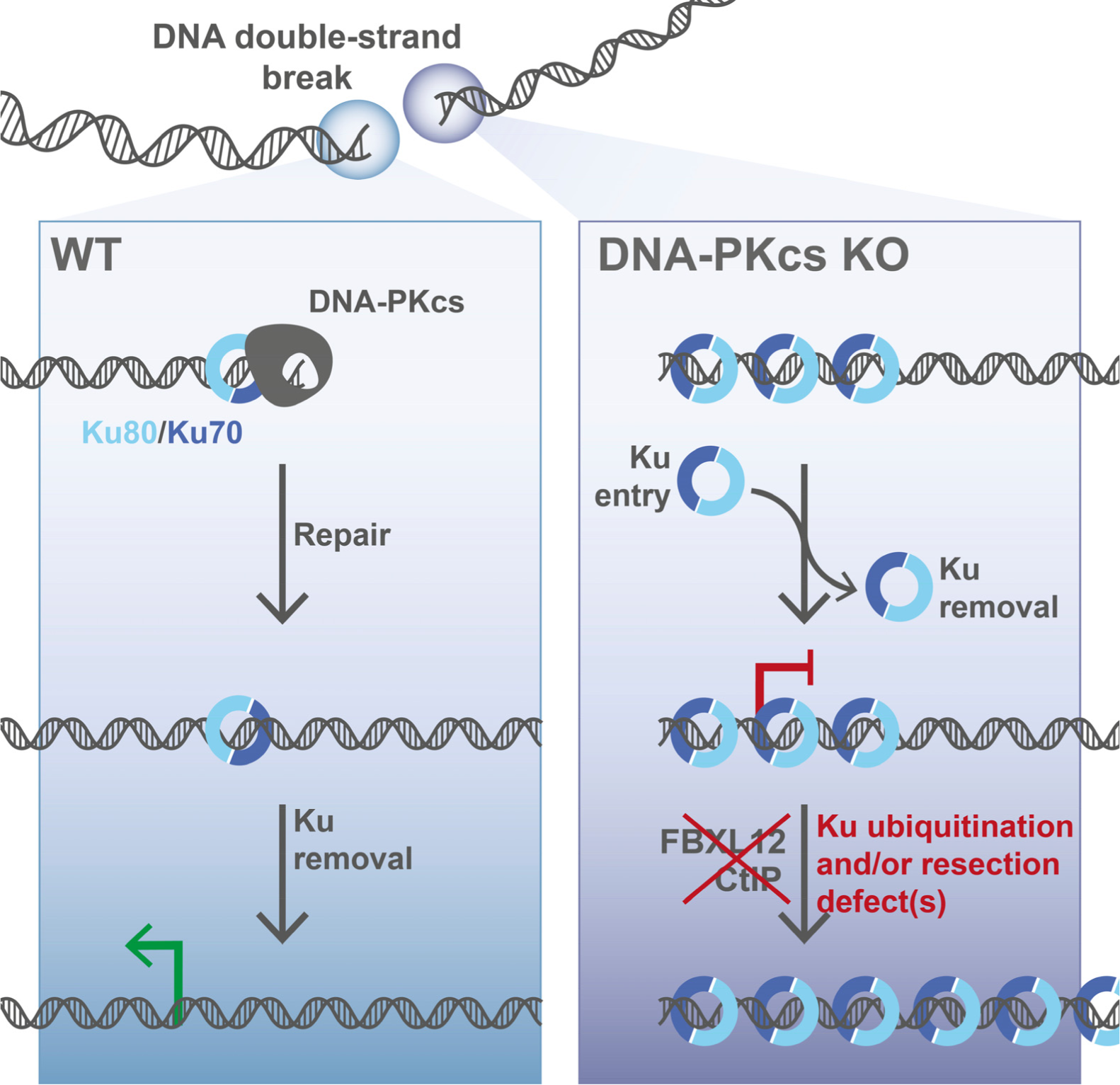

## Introduction

The DNA of living organisms is continuously exposed to endogenous and exogenous DNA damaging agents which induce a variety of DNA lesions (Ciccia and Elledge, 2010). Among these lesions, DNA double-strand breaks (DSBs) are particularly toxic. Recognition of DSBs activates the DNA damage response (DDR) — an integrated response that blocks cell cycle progression, activates a transcriptional program, and attempts to repair the lesion through several DNA repair mechanisms (Ciccia and Elledge, 2010). In vertebrates, DSBs are recognized by two main sensors: the MRE11-RAD50-NBS1 complex (MRN) and the Ku complex, a ring-shaped heterodimer of Ku70-Ku80. MRN clamps on the flank of the DNA end, recruits the ATM protein kinase and possesses nuclease activities involved in the initiation of DNA end resection. In contrast, Ku accommodates one DNA end in its central cavity and protects it from unscheduled exonucleolytic processing (Liang and Jasin, 1996; Walker et al., 2001). In addition to detecting and binding free DNA ends, Ku initiates Non-Homologous End Joining (NHEJ), the main DSB repair pathway for two-ended DSBs, which tethers and ligates the two DNA ends together. Ku promotes NHEJ by serving as an assembly site for most of the NHEJ proteins, including the DNA-dependent protein kinase catalytic subunit (DNA-PKcs), the XRCC4-DNA ligase IV ligation complex, and the scaffold proteins XLF/Cernunnos and PAXX (Frit et al., 2019; Stinson and Loparo, 2021). The DNA:Ku:DNA-PKcs complex forms the active DNA-PK kinase that acts on numerous substrates, including itself, and is essential for the NHEJ-mediated repair of DNA ends which require processing (Chang et al., 2016; Cisneros-Aguirre et al., 2022; Stinson et al., 2020).

Ku is an abundant nuclear protein (Carter et al., 1990), with high affinity for DNA ends (binding constant of 2.4 nM (Blier et al., 1993)). *In vitro* multiple Ku molecules thread onto DNA from free ends with the amount of Ku loading only limited by the size of the DNA fragment and the amount of available Ku (Blier et al., 1993; de Vries et al., 1989). However, in cells an average of only 2-3 Ku molecules are found at DSBs (∼1 per end, (Britton et al., 2013)), indicating that there are mechanisms that prevent multiple Ku proteins from loading onto DNA. Preventing Ku entry into chromatin may be important to allow for other genome transactions. In agreement, Ku overloading *in vitro* blocks the association of transcription factors to their binding sites (e.g. Sp1, AP1, Oct1/OTF1 (Ono et al., 1996)), reduces their transcription (Frit et al., 2000) and impairs repair of other nearby lesions by dedicated mechanisms, such as Nucleotide Excision Repair (Frit et al., 1998; Frit et al., 2000).

The stoichiometry of Ku at DNA ends may be regulated by a combination of passive and active mechanisms. Ku is known to interact with many DDR proteins and these factors may sterically block additional Ku binding or recruit proteins that remove Ku from DNA ends. Prior work showed that inhibiting DNA-PKcs attenuates Ku loading within human cell extracts suggesting that it may function as a regulatable barrier (Calsou et al., 1999). This raises the question of the role of the other NHEJ factors in blocking Ku entry into chromatin. In addition, a number of active mechanisms promote Ku eviction from DNA ends and chromatin. First, the nucleolytic activities of the CtIP-MRN complex, stimulated by ATM-dependent CtIP phosphorylation, antagonizes the persistence of Ku at single-ended DSBs generated in S-phase by the DNA topoisomerase I poison camptothecin (CTP, (Chanut et al., 2016)). This mechanism requires the phosphorylation of the DNA-PKcs ABCDE sites by ATM and its dissociation from the NHEJ complex prior to Ku release (Britton et al., 2020). Second, the splicing factor XAB2 antagonizes Ku binding at DSBs produced by CPT and temozolomide by a mechanism which remains to be characterized (Sharma et al., 2021). Third, direct Ku phosphorylation promotes its release in S-phase to favor other DNA repair pathways (Lee et al., 2016). Finally, Ku undergoes polyubiquitination and release from ends by the activity of the unfoldase VCP (p97 or Cdc48) (Postow et al., 2008; van den Boom et al., 2016). Several E3 ubiquitin ligases, including RNF8, RNF138, RNF126 and FBXL12, have been implicated in promoting Ku release through direct ubiquitination (Fowler et al., 2022; Ishida et al., 2017; Ismail et al., 2015; Kolas et al., 2007; Postow and Funabiki, 2013). Importantly, Ku release was shown to be largely dependent on neddylation, *i.e.* the conjugation of the ubiquitin-like NEDD8 to Cullins, implying that a Cullin-based ubiquitin ligase is the key Ku releasing factor (Brown et al., 2015).

Here we investigate the cellular mechanisms regulating Ku stoichiometry on DNA using both human cells and *Xenopus* egg extracts. Using two complementary super-resolution methods to analyze Ku foci at individual DSBs in human cells, we reveal that DNA-PKcs is a major barrier which prevents multiple Ku molecules from loading onto chromatin. Using a quantitative single-molecule assay within *Xenopus* egg extracts to measure the precise amount of Ku at DNA ends, we establish in another vertebrate that DNA-PKcs plays a key structural role in enforcing the 1:1 Ku:DNA end stoichiometry. We also demonstrate that without DNA-PKcs a neddylation/FBXL12-dependent process actively prevents excessive Ku accumulation into chromatin, most likely through direct Ku ubiquitination. In addition, we reveal that a CtIP/ATM-dependent process quickly reverses aberrant Ku accumulation on chromatin during S-phase, which suggests that excess Ku can also be actively removed by DNA end resection. Finally, through developing an assay to monitor in cells how Ku entry onto DNA impacts transcription in the vicinity of DNA ends, we demonstrate that DNA-PKcs is essential to protect transcription from inhibition by Ku overloading. Altogether this work identifies DNA-PKcs as the main barrier that prevents Ku overloading onto chromatin and demonstrates that additional mechanisms act to remove excess Ku molecules when needed.

## Results

### DNA-PKcs physically limits Ku entry into chromatin in human cells

To determine the impact of DNA ligase IV (Lig4) and DNA-PKcs on Ku loading at individual DSBs, we used structured illumination super-resolution microscopy (SIM) to measure the intensity of Ku foci following ionizing radiation (IR) in wild-type (WT) U2OS cells, U2OS cells knocked-out for DNA-PKcs (PKcs-KO), DNA ligase IV (LIG4-KO) or both (LIG4/PKcs-KO, **Fig.1A**). These cells lines displayed the expected sensitivity to IR (**Fig.S1A**). As soon as five min after IR, Ku foci were on average 2-3 fold brighter in cells lacking DNA-PKcs and this increase persisted for one hour after irradiation (**Fig.1B**). Loss of Lig4, either alone or in combination with DNA-PKcs, had no measurable impact on Ku foci, suggesting that the NHEJ defect is not responsible for the Ku accumulation observed on irradiated chromatin in PKcs-KO cells. Next, we examined the role of DNA-PKcs kinase activity on Ku foci intensity. Cells were pre-treated with the specific DNA-PK inhibitor nedisertib (PKi, aka M3814 (Zenke et al., 2020)) and then the same analysis was performed one hour after IR in PKcs-KO or WT cells. Ku foci were brighter in PKcs-KO cells in contrast to WT cells treated with PKi (**Fig.1C**), supporting a physical rather than a catalytic role of DNA-PKcs in limiting Ku loading at DSBs.

**Figure 1:**
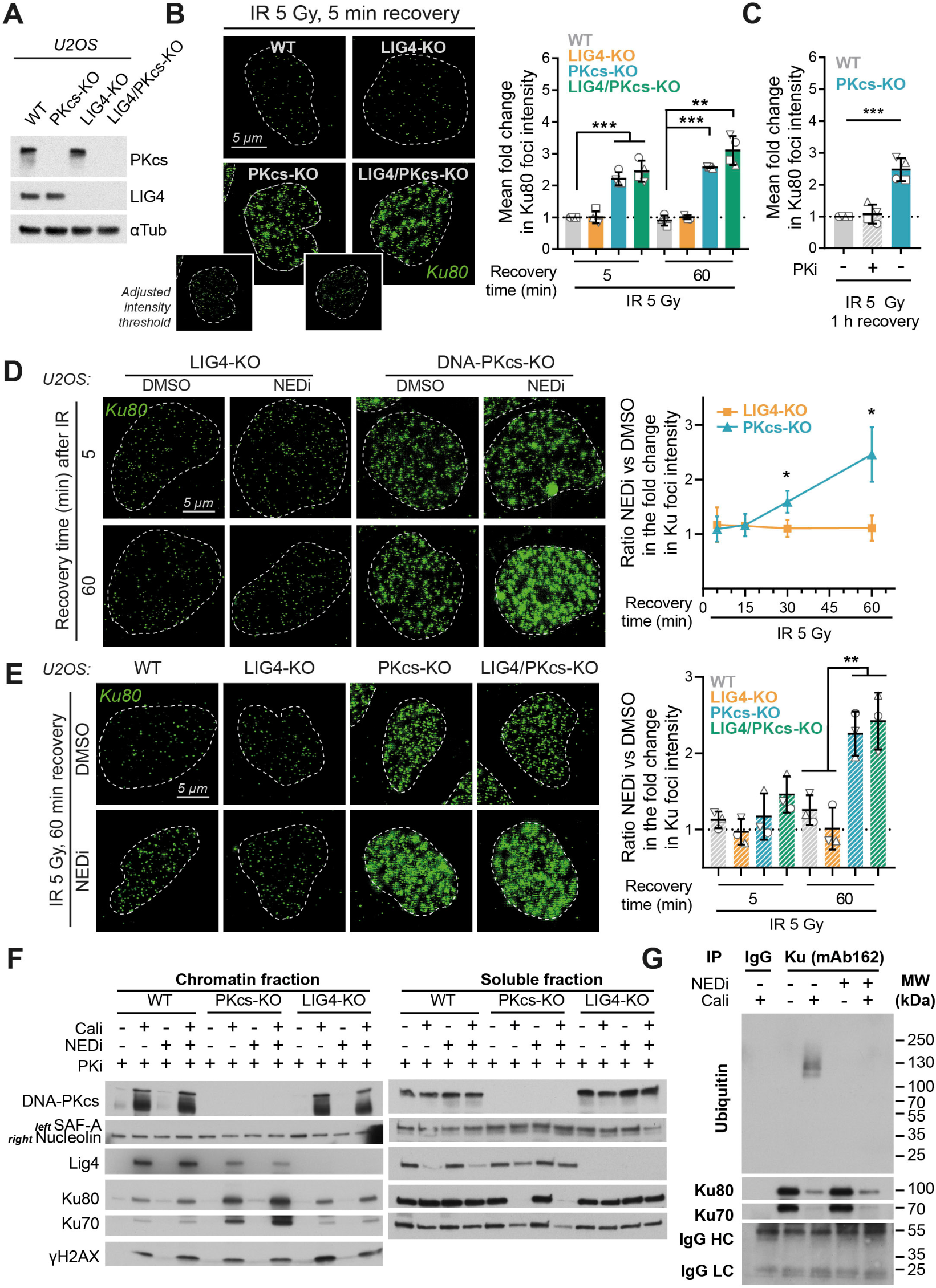
DNA-PKcs presence but not activity limits Ku accumulation at DSBs. **A.** Immunoblot of U2OS cells wild-type (WT) or knocked-out for the indicated genes. See **Fig.S1A** for the IR sensitivity analysis of each of these cells. **B.** U2OS WT, DNA-PKcs KO (PKcs-KO), DNA ligase IV KO (LIG4-KO) or DNA-PKcs and Lig4 KO (LIG4/PKcs-KO) received 5 Gy of IR before being post-incubated 5 or 60 min and processed for Ku foci imaging. Representative pictures are shown on the left panel. Ku foci average intensity was measured and normalized to the Ku foci intensity measured after 5 Gy of IR in WT U2OS to compute the fold change in Ku foci intensity in each condition, depicted on the graph on the right panel. **C.** WT or PKcs-KO U2OS cells received 5 Gy of IR and were post-incubated for 60 min before being processed for Ku foci imaging. Cells were pre-treated with 250 nM nedisertib (PKi) for 1 h before treatment when indicated. Fold change in Ku foci intensity in each condition is displayed. **D.** PKcs-KO or LIG4-KO U2OS cells received 5 Gy of IR and were post-incubated for the indicated time with or without NEDi before being processed for Ku foci imaging. Representative pictures are shown on the left, while the ratio between the changes of Ku foci intensity with NEDi versus without NEDi (DMSO) are plotted on the graph on right. The corresponding graph depicting the fold change in Ku foci intensity in each condition corresponds to **Fig.S1B**. **E.** WT, PKcs-KO, LIG4-KO or LIG4/PKcs-KO U2OS cells received 5 Gy of IR and were post-incubated 5 or 60 min with or without NEDi before being processed for Ku foci imaging. Representative pictures are shown on the left panel, while the ratio between the change of Ku foci intensity with NEDi versus without NEDi (DMSO) is plotted on the graph on right. The corresponding graph depicting the fold change in Ku foci intensity in each condition corresponds to **Fig.S1C**. **Fig. S1D,E** and **Fig.S1F,G** depict similar analyses in HeLa cells and in complemented PKcs-KO U2OS, respectively, while **Fig.S2** reports the use of STORM imaging to monitor the size and shape of Ku foci in absence of DNA-PKcs +/-NEDi. **F.** WT, PKcs-KO or LIG4-KO U2OS incubated with 3 µM of NU7441 (PKi) were treated with 3 nM Cali for 1 h with or without NEDi, before being collected and processed to separate a chromatin fraction from a soluble fraction which were both analyzed by immunoblotting. SAF-A and nucleolin were used as loading controls for the chromatin and the soluble fraction, respectively. **G.** Control (IgG) or anti-Ku immunoprecipitation were performed from the soluble fractions to monitor Ku ubiquitination in response to Cali with or without NEDi.

Given that a neddylation-dependent process is the main driver of Ku removal from chromatin after completion of DNA repair (Brown et al., 2015), we monitored the impact of the neddylation inhibitor MLN4924 (NEDi, (Soucy et al., 2009)) on Ku accumulation at DSBs. In PKcs-KO cells, inhibition of neddylation led to a time-dependent increase in Ku foci intensity up to ∼2.5-fold 1 h after IR relative to PKcs-KO without NEDi (**Fig.1D,S1B**), suggesting that a neddylation-dependent machinery actively operates in these cells to remove excess Ku from chromatin. No effect of NEDi was observed in the absence of DNA ligase IV alone (**Fig.1E,S1C**), supporting the model that the observed increase in Ku foci intensity is not the result of a DNA repair defect, but rather linked to the physical function of DNA-PKcs in antagonizing Ku entry into chromatin. Similar results were obtained in HeLa cells (**Fig. S1D,E**): Ku foci were ∼2 times brighter in HeLa PKcs-KO as compared to HeLa WT both at 5 min and 60 min after IR. Inhibition of neddylation further increases the intensity of Ku foci in HeLa PKcs-KO, which reached ∼4 times the intensity of Ku foci in WT cells, while it had nearly no effect on HeLa WT cells. Our findings are further validated through rescue experiments conducted in U2OS PKcs-KO (**Fig.S1F,G**): in U2OS PKcs-KO complemented with an empty plasmid, Ku foci are brighter both at 5 min and 60 min after IR as compared to U2OS complemented with WT DNA-PKcs, and with inhibition of neddylation an additional increase in Ku foci intensity is observed 60 min after IR. In contrast, Ku foci intensity was similar to U2OS WT cells when U2OS PKcs-KO cells were complemented with a DNA-PKcs WT expressing plasmid. Inhibiting DNA-PK activity with another well-characterized catalytic inhibitor (NU7441), did not lead to an increase in Ku foci intensity in WT cells (**Fig.S1G**).

Since the resolution of our 3D-SIM imaging set-up is ∼150 nM, we adapted our imaging protocol to STORM with a theoretical resolution of ∼50 nm (20 nm plus the ∼30 nm resolution lost as a result of using primary-secondary immunocomplexes for detection (Jimenez et al., 2020)). While the sizes of the longest side of a single Ku molecule (∼10 nm) or of two Ku molecules (∼21 nm) in the long range synaptic complex (Seif-El-Dahan et al., 2023) are below the resolution of our setup, we hypothesized that, under conditions of Ku overloading, STORM imaging may reveal differences in foci size and organization not accessible to our 3D-SIM imaging approach. For this purpose, Ku foci were analyzed 5 min after IR in WT cells or 1 h after IR in NHEJ-deficient cells (LIG4-KO), co-inactivated or not for DNA-PKcs (LIG4/PKcs KO). When indicated the cells were also treated with NEDi. To identify Ku foci from the coordinates of localized molecules obtained by STORM, we used a previously described approach based on Voronoï tessellation (Levet et al., 2015; Levet and Sibarita, 2023)). Our analysis shows that the loss of DNA-PKcs leads to larger Ku foci, with a mean size of ∼67 nm, in contrast to WT and LIG4-KO conditions in which the mean size is ∼60 nm (**Fig.S2A**). Ku foci size in PKcs-KO was further increased with NEDi (mean size ∼83 nm). In line with an increase in the size of Ku foci, the number of localizations per cluster was also increased in the absence of DNA-PKcs and further exacerbated upon NEDi treatment (**Fig.S2B**), supporting the fact that a higher number of Ku molecules is present at DSB sites in the absence of DNA-PKcs and that this level is further enhanced when neddylation is blocked. Ku foci in the absence of DNA-PKcs have a lower density than in the WT condition, both without and with NEDi (**Fig.S2C**), and form elongated structures (**Fig.S2D,E**), suggesting the spreading of Ku molecules along the DNA. Altogether, these data provide additional evidence that, upon loss of DNA-PKcs, Ku invades the chromatin on the flank of each DSB.

To complement our observations on the mechanisms controlling Ku hyper accumulation at DSBs, we used an orthogonal biochemical fractionation assay (Drouet et al., 2005) to monitor the association of NHEJ proteins to chromatin in response to a radiomimetic drug, calicheamicin-γ1 (Cali, (Zein et al., 1988)). WT, PKcs-KO and LIG4-KO U2OS cell lines were pretreated with PKi to equally neutralize their NHEJ capacities (**Fig.1F**). Similar to our SIM imaging results, loss of DNA-PKcs led to a strong increase in Ku association to chromatin upon Cali treatment as compared to WT and LIG4-KO cells, even resulting in Ku partial depletion from the soluble protein fraction (**Fig.1F**). Of note, Lig4 association to damaged chromatin was decreased in PKcs-KO cells, reflecting the stabilization of Lig4 at DSBs mediated by DNA-PKcs (Calsou et al., 2003; Drouet et al., 2005). Addition of NEDi to the cells resulted in a further increase in the amount of Ku associated with chromatin only in the absence of DNA-PKcs, highlighting the role for neddylation in limiting Ku accumulation onto chromatin. Notably, when Ku was immunoprecipitated from the soluble fraction from PKcs-KO cells, a ubiquitination signal was detected above the Ku80 band after Cali treatment, which disappeared with NEDi (**Fig.1G**), supporting the involvement of a Cullin-based ubiquitin ligase in promoting Ku release from the damaged chromatin through direct Ku ubiquitination.

### DNA-PKcs controls Ku stoichiometry at DSB ends

To quantitatively assess the function of DNA-PKcs in regulating the Ku stoichiometry at DSBs, we developed a single molecule assay in *Xenopus* egg extracts to measure the number of fluorescently labeled Ku molecules loaded onto surface-attached DNA substrates (**Fig.2A**). Ku was visualized by complementing Ku-immunodepleted extracts with purified *Xenopus laevis* Halo-Ku80:Ku70 (**Fig.S3A,B**), which retained wild-type activity in NHEJ assays (**Fig.S3C**). To determine the maximum number of Ku molecules which can load onto a 100 base pairs Cy3-labeled DNA substrate tethered to the surface of a microfluidic flowcell, we first incubated Cy5-labeled Halo-Ku80:Ku70 with the tethered DNA for 60 min in wash buffer. Next, we imaged both the surface-bound DNA and Ku using total internal reflection fluorescence imaging (**Fig. 2A**) and used the number of photobleaching steps as a readout of the number of Ku molecules on the DNA substrate (**Fig.2B**). Photobleaching analysis of colocalized Ku foci found that multiple Ku molecules would load onto individual substrates with a mean of 3 ± 2.2 Ku molecules per DNA (**Fig. 2C**, see for each condition **Fig.S3G,H,I** and **Fig.S3J** corresponding respectively to single-molecule probability distributions and representative trajectories). In contrast, when tethered DNA was incubated with *Xenopus* egg extracts in which endogenous Ku was replaced by Halo-Ku80:Ku70 (**Fig.S3B**), only 1 ± 0.4 Ku molecules loaded per DNA substrate (**Fig.2D**), showing that a mechanism exists to prevent multiple Ku molecules threading onto DNA under physiological conditions. Notably, immunodepletion of DNA-PKcs from the egg extract (**Fig.S3D**) resulted in overloading of Ku molecules (2 ± 1.5) (**Fig.2E**), consistent with DNA-PKcs acting as a critical regulator of Ku loading. In contrast, immunodepletion of the core factors XLF and XRCC4-LIG4 from egg extract (**Fig.S3E**) had no significant impact on Ku stoichiometry (1 ± 0.5) (**Fig.2F**).

**Figure 2:**
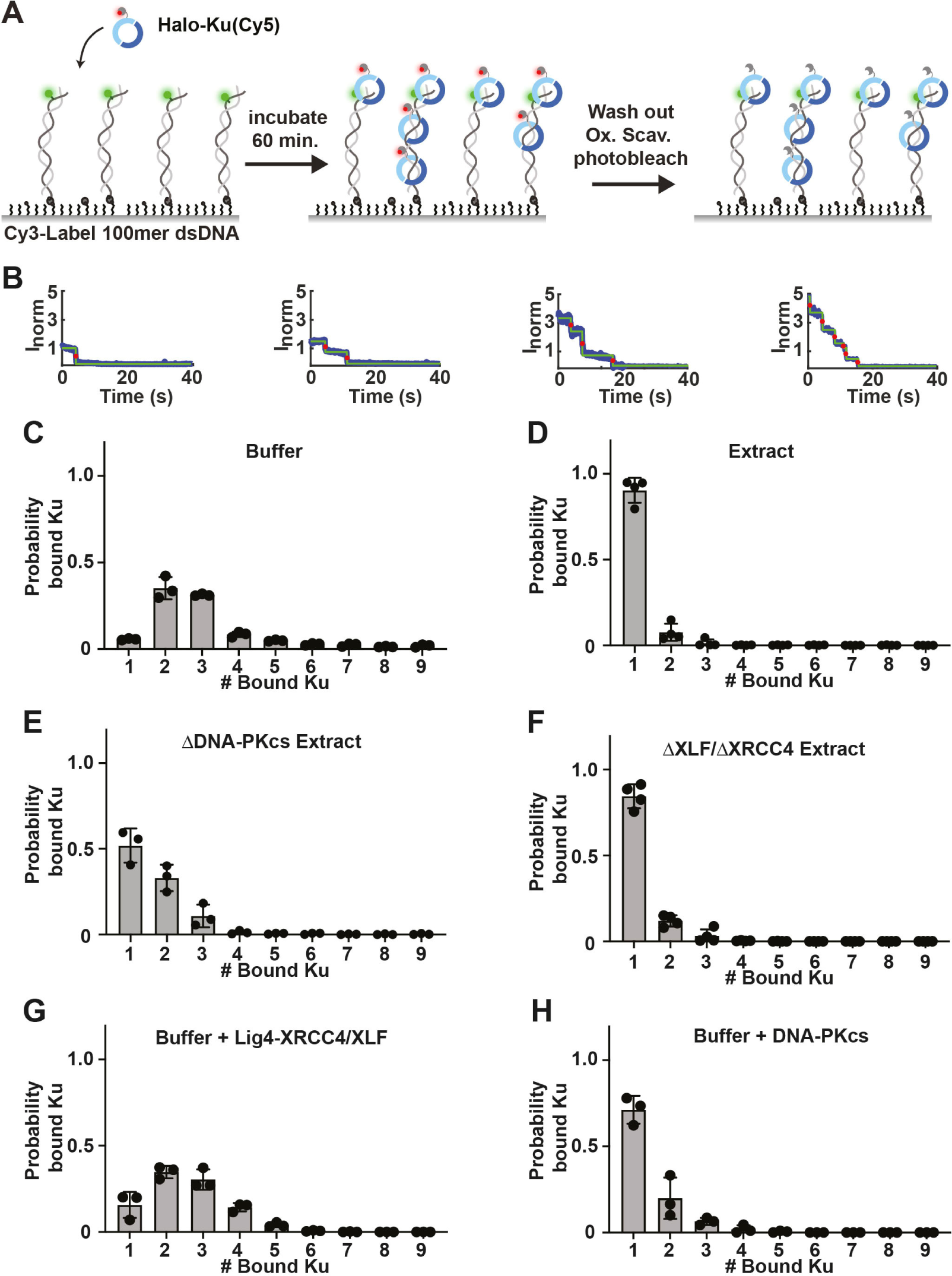
DNA-PKcs prevents Ku overloading onto a single DNA end. **A.** Scheme depicting the single molecule assay used to quantify Ku loading on DNA in presence or in absence of DNA-PKcs. A Cy3-labeled 100 bp DNA substrate was attached to a glass surface, incubated for 60 min with Cy5-labelled xenopus Halo-Ku80:Ku70 alone or in non-cycling *Xenopus* eggs extracts. After incubation and wash-out of the oxygen scavenger, the photobleaching of individual Ku molecules was monitored under continuous illumination and the number of bleaching steps was used as a readout of the number of Ku molecules on DNA. **B.** Representative trajectories highlighting photobleaching events for Halo-Ku80:Ku70 bound to DNA in buffer. Representative trajectories for each condition are shown in **Fig.S3J**. **C-H**: On each panel the normalized histograms depicting fractional occupancy of Ku70/80 on DNA ends, constructed from mean fractions per occupancy bin calculated in 3 independent experiments. The number of events and total number of molecules observed for each experiment are reported in **Table 1**. **C.** The number of Ku molecules was monitored as described in **A.** using purified Cy5-labeled Ku incubated in ELB wash buffer. **D.** The number of Ku molecules was monitored as described in **A.** using *Xenopus* eggs extracts containing Cy5-labeled Ku. **E.** The number of Ku molecules was monitored as described in **A.** using *Xenopus* eggs extracts immunodepleted for DNA-PKcs (ΔDNA-PKcs) and containing purified Cy5-labeled Ku. **F.** The number of Ku molecules was monitored as described in **A.** using *Xenopus* eggs extracts immunodepleted for XLF and XRCC4 (ΔXLF/ΔXRCC4) and containing purified Cy5-labeled Ku. **G.** The number of Ku molecules was monitored as described in **A.** using purified Cy5-labeled Ku mixed with 60 nM Lig4-XRCC4, 60 nM XLF, and incubated in ELB wash buffer. **H.** The number of Ku molecules was monitored as described in **A.** using purified Cy5-labeled Ku mixed with 60 nM DNA-PKcs and incubated in ELB wash buffer.

**Table 1.** Single-molecule Photobleaching events.

| **Ku bound to 100 mer in buffer** |  |  |  |  |  |  |  |  |  |  |  |  |  |
| --- | --- | --- | --- | --- | --- | --- | --- | --- | --- | --- | --- | --- | --- |
| Replicate | Tethered DNA-Ku complex (#) | Mean bound Ku (#) | Stdev (#) | (#) DNA bound by: |  |  |  |  |  |  |  |  |  |
|  |  |  |  | 1 Ku | 2 Ku | 3 Ku | 4 Ku | 5 Ku | 6 Ku | 7 Ku | 8 Ku | 9 Ku | >=10 Ku |
| A | 1321 | 3 | 1.9 | 82 | 573 | 419 | 89 | 60 | 28 | 19 | 16 | 3 | 35 |
| B | 1196 | 4 | 2.4 | 67 | 379 | 405 | 129 | 59 | 39 | 34 | 20 | 17 | 64 |
| C | 1334 | 3 | 2.2 | 76 | 466 | 436 | 126 | 75 | 41 | 41 | 15 | 17 | 58 |
| **Ku bound to 100 mer in buffer with 60 nM Lig4-XRCC4 and 60 nM XLF** |  |  |  |  |  |  |  |  |  |  |  |  |  |
| Replicate | Tethered DNA-Ku complex (#) | mean bound Ku (#) | Stdev (#) | (#) DNA bound by: |  |  |  |  |  |  |  |  |  |
|  |  |  |  | 1 Ku | 2 Ku | 3 Ku | 4 Ku | 5 Ku | 6 Ku | 7 Ku | 8 Ku | 9 Ku | >=10 Ku |
| A | 987 | 3 | 1.0 | 42 | 216 | 223 | 95 | 21 | 2 | 0 | 0 | 0 | 6 |
| B | 775 | 3 | 1.2 | 127 | 297 | 155 | 133 | 57 | 6 | 0 | 0 | 0 | 6 |
| C | 952 | 2 | 1.1 | 192 | 356 | 257 | 110 | 31 | 6 | 0 | 0 | 0 | 6 |
| **Ku bound to 100 mer in buffer with 60 nM DNA-PKcs** |  |  |  |  |  |  |  |  |  |  |  |  |  |
| Replicate | Tethered DNA-Ku complex (#) | mean bound Ku (#) | Stdev (#) | (#) DNA bound by: |  |  |  |  |  |  |  |  |  |
|  |  |  |  | 1 Ku | 2 Ku | 3 Ku | 4 Ku | 5 Ku | 6 Ku | 7 Ku | 8 Ku | 9 Ku | >=10 Ku |
| A | 4468 | 1 | 0.6 | 622 | 332 | 44 | 1 | 0 | 0 | 0 | 0 | 0 | 4 |
| B | 309 | 1 | 0.9 | 241 | 31 | 20 | 14 | 3 | 0 | 0 | 0 | 0 | 5 |
| C | 1081 | 1 | 0.7 | 286 | 64 | 33 | 4 | 2 | 0 | 0 | 0 | 0 | 5 |
| **Ku bound to 100 mer in *Xenopus* Egg Extract** |  |  |  |  |  |  |  |  |  |  |  |  |  |
| Replicate | Tethered DNA-Ku complex (#) | mean bound Ku (#) | Stdev (#) | (#) DNA bound by: |  |  |  |  |  |  |  |  |  |
|  |  |  |  | 1 Ku | 2 Ku | 3 Ku | 4 Ku | 5 Ku | 6 Ku | 7 Ku | 8 Ku | 9 Ku | >=10 Ku |
| A | 350 | 1 | 0.5 | 323 | 23 | 1 | 1 | 1 | 1 | 0 | 0 | 0 | 0 |
| B | 309 | 1 | 0.7 | 246 | 46 | 14 | 1 | 0 | 1 | 0 | 1 | 0 | 0 |
| C | 809 | 1 | 0.2 | 766 | 41 | 2 | 0 | 0 | 0 | 0 | 0 | 0 | 0 |
| **Ku bound to 100 mer in *Xenopus* Egg Extract ΔXLF/ΔXRCC4** |  |  |  |  |  |  |  |  |  |  |  |  |  |
| Replicate | Tethered DNA-Ku complex (#) | mean bound Ku (#) | Stdev (#) | (#) DNA bound by: |  |  |  |  |  |  |  |  |  |
|  |  |  |  | 1 Ku | 2 Ku | 3 Ku | 4 Ku | 5 Ku | 6 Ku | 7 Ku | 8 Ku | 9 Ku | >=10 Ku |
| A | 466 | 1 | 0.4 | 394 | 68 | 3 | 1 | 0 | 0 | 0 | 0 | 0 | 0 |
| B | 436 | 1 | 0.5 | 351 | 67 | 17 | 1 | 0 | 0 | 0 | 0 | 0 | 0 |
| C | 515 | 1 | 0.5 | 429 | 69 | 13 | 4 | 0 | 0 | 0 | 0 | 0 | 0 |
| **Ku bound to 100mer in *Xenopus* Egg Extract ΔDNA-PKcs** |  |  |  |  |  |  |  |  |  |  |  |  |  |
| Replicate | Tethered DNA-Ku complex (#) | mean bound Ku (#) | Stdev (#) | (#) DNA bound by: |  |  |  |  |  |  |  |  |  |
|  |  |  |  | 1 Ku | 2 Ku | 3 Ku | 4 Ku | 5 Ku | 6 Ku | 7 Ku | 8 Ku | 9 Ku | >=10 Ku |
| A | 1082 | 2 | 1.5 | 676 | 301 | 64 | 22 | 2 | 6 | 2 | 2 | 2 | 7 |
| B | 1046 | 2 | 1.5 | 689 | 260 | 53 | 15 | 10 | 7 | 5 | 3 | 3 | 4 |
| C | 2087 | 2 | 1.5 | 848 | 836 | 381 | 9 | 5 | 2 | 1 | 0 | 1 | 5 |
| **Ku bound to 100 mer in *Xenopus* Egg Extract + NEDi** |  |  |  |  |  |  |  |  |  |  |  |  |  |
| Replicate | Tethered DNA-Ku complex (#) | mean bound Ku (#) | Stdev (#) | (#) DNA bound by: |  |  |  |  |  |  |  |  |  |
|  |  |  |  | 1 Ku | 2 Ku | 3 Ku | 4 Ku | 5 Ku | 6 Ku | 7 Ku | 8 Ku | 9 Ku | >=10 Ku |
| A | 342 | 1 | 0.9 | 301 | 34 | 1 | 1 | 2 | 0 | 2 | 0 | 0 | 1 |
| B | 841 | 1 | 1.4 | 675 | 142 | 16 | 0 | 4 | 1 | 1 | 1 | 0 | 1 |
| C | 1121 | 1 | 0.8 | 1055 | 60 | 4 | 1 | 0 | 0 | 0 | 0 | 0 | 1 |
| **Ku bound to 100 mer in *Xenopus* Egg Extract ΔDNA-PKcs + NEDi** |  |  |  |  |  |  |  |  |  |  |  |  |  |
| Replicate | Tethered DNA-Ku complex (#) | mean bound Ku (#) | Stdev (#) | (#) DNA bound by: |  |  |  |  |  |  |  |  |  |
|  |  |  |  | 1 Ku | 2 Ku | 3 Ku | 4 Ku | 5 Ku | 6 Ku | 7 Ku | 8 Ku | 9 Ku | >=10 Ku |
| A | 1462 | 2 | 1.2 | 815 | 504 | 82 | 17 | 13 | 12 | 6 | 6 | 1 | 7 |
| B | 1453 | 2 | 1.2 | 861 | 376 | 136 | 40 | 13 | 12 | 3 | 0 | 6 | 12 |
| C | 1899 | 2 | 1.2 | 539 | 919 | 367 | 43 | 11 | 7 | 3 | 2 | 3 | 8 |

DNA-PKcs may play a direct structural role in limiting Ku stoichiometry by retaining Ku at the DNA end and sterically blocking additional Ku molecules from loading. Alternatively, DNA-PKcs may recruit additional factors that regulate Ku binding. To distinguish between these two scenarios, we preincubated fluorescently labeled Ku with purified *Xenopus* NHEJ factors and then added them to the flow cell. The number of DNA bound Ku molecules were then counted after 60 min of incubation. While the addition of XLF and XRCC4-LIG4 had no noticeable effect on Ku loading compared to the Ku only control (3 ± 1.2) (**Fig.2G**), premixing Ku and DNA-PKcs purified from extract (**Fig.S3F**) led to a substantial reduction in Ku stoichiometry (1 ± 0.6) (**Fig.2H**) which mimicked observations of undepleted egg extract. These results demonstrate that DNA-PKcs acts as a structural barrier to Ku loading that is independent of its kinase activity.

Next, we examined if active removal of Ku via ubiquitination contributes to limiting Ku stoichiometry in egg extract. To determine if Ku was undergoing polyubiquitination under our assay conditions, we bound the 100 bp DNA substrate to streptavidin magnetic beads and incubated with extracts to monitor association of the DNA-PK complex (**Fig.S4A**). Ku and DNA-PKcs rapidly associated with DNA ends while a slower migrating Ku80 species appeared over time that was sensitive to treatment with the deubiquitinase (DUB) USP2 (**Fig.S4B**). Addition of the neddylation inhibitor NEDi MLN4924, which did not impact end joining (**Fig.S4C**), resulted in loss of these modified Ku species and stabilized Ku at DNA ends (**Fig.S4B**). Together these results demonstrate that a neddylation-dependent polyubiquitinated form of Ku80 is generated in our experiments. However, our single-molecule imaging revealed that the addition of NEDi did not significantly alter Ku stoichiometry in the presence (1 ± 1.1 Ku molecules per DNA) (**Fig.S4D**) or absence of DNA-PKcs (2 ± 1.2) (**Fig.S4E**) as compared to the relevant conditions in the absence of inhibitor (see **Fig.S4F,G,H** and **Fig.S4I** respectively for single-molecule probability distributions and representative trajectories). Collectively, these results provide further evidence that DNA-PKcs plays a prominent role in limiting Ku entry onto DNA in vertebrates. Importantly, this regulation of Ku stoichiometry appears unique to DNA-PKcs as loss of the core factors XLF and XRCC4-LIG4 had no significant impact on Ku stoichiometry.

### In S-phase, an ATM-dependent mechanism overcomes Ku accumulation

Multiple mechanisms function in S-phase to antagonize Ku loading on DSBs, with ATM playing a prominent role in controlling these mechanisms (Britton et al., 2020; Chanut et al., 2016; Sharma et al., 2021). To evaluate the relative contribution of neddylation and ATM in regulating Ku entry into chromatin in and outside of S-phase, we used the specific ATM inhibitor KU-55933 (ATMi) alone or in combination with NEDi and monitored Ku accumulation at DSBs post IR in PKcs-KO cells in which Ku overloading is expected. PCNA co-staining was used to identify cells in S-phase. DNA-PKcs KO led to the expected Ku overloading 5 min after ionizing radiation in both replicating and non-replicating cells. However, the mean Ku foci intensity decreased over time in PCNA positive cells (**Fig.3A**), suggesting an S-phase specific mechanism for Ku removal from chromatin. Inhibiting neddylation led to a time-dependent increase in Ku loading both in non-replicating and replicating cells. However, in replicating cells, Ku foci intensity was ∼2-times lower at 4 h compared to non-replicating cells, implying that a Ku release mechanism operates in S-phase independently of the neddylation-dependent removal from damaged chromatin. Notably, inhibiting ATM prevented the preferential removal of Ku in S-phase, either with or without NEDi, suggesting that ATM controls Ku eviction from chromatin in S-phase. This supports that two active mechanisms operate to limit Ku accumulation in DNA-PKcs-deficient cells: a neddylation-dependent mechanism operating throughout the cell cycle and an ATM-dependent mechanism operating in S-phase. We therefore attempted to provide additional insights into these two mechanisms removing Ku from chromatin when DNA-PKcs is absent.

**Figure 3:**
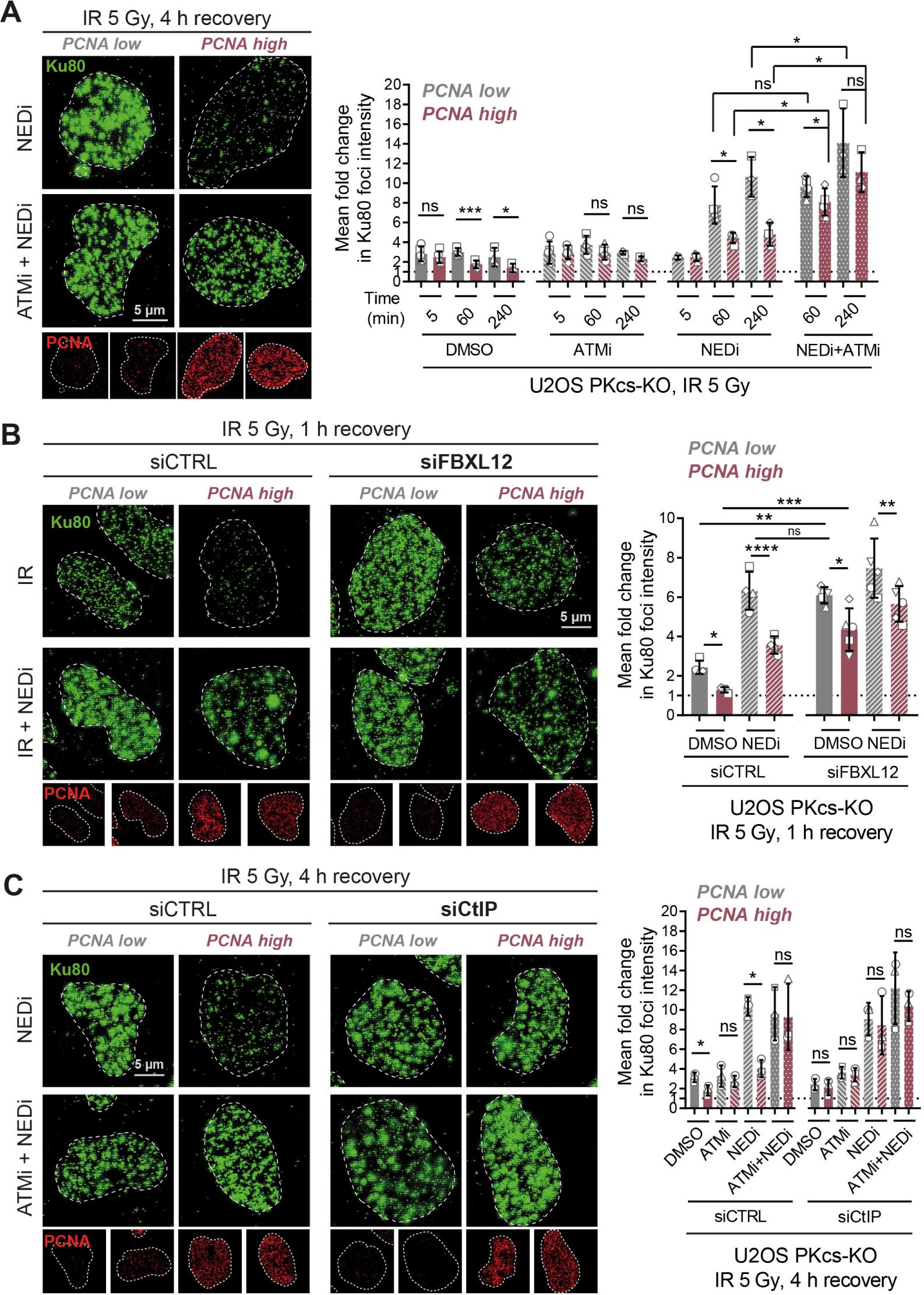
Different mechanisms limit Ku loading during the cell cycle. **A.** PKcs-KO U2OS were pre-treated with NEDi, ATMi or both and received 5 Gy of IR before being post-incubated for the indicated time before being processed for immunofluorescence. A PCNA staining was used to identify the cells in S-phase. Representative pictures are shown on the left panel, with insets at the bottom to illustrate the PCNA staining. Ku foci average intensity was measured and normalized to the Ku foci intensity in U2OS WT measured 5 min after 5 Gy of IR to compute the fold change in Ku foci intensity in each condition, displayed on the graph on the right panel. **B.** PKcs-KO U2OS were transfected by siRNA control or against FBXL12 before being treated and processed as described in A. Representative pictures are shown on the left panel, with insets at the bottom to illustrate the PCNA staining. An immunoblot showing the depletion of FBXL12 is shown in **Fig.S5A**, while the graph on the right panel shows the fold change in Ku foci intensity computed as in A. **C.** PKcs-KO U2OS were transfected by siRNA control or against CtIP before being treated and processed as described in A. Representative pictures are shown on the left panel, with insets at the bottom to illustrate the PCNA staining. An immunoblot showing the depletion of CtIP is shown in **Fig.S5E**, while the graph on the right panel shows the fold change in Ku foci intensity computed as in A.

### FBXL12 mediates the neddylation-dependent removal of Ku from chromatin

Given that FBXL12 is a Cullin-dependent E3 ubiquitin ligase implicated in Ku removal after repair (Fowler et al., 2022; Postow and Funabiki, 2013), we used siRNA-mediated depletion to test its role in the neddylation-dependent removal of Ku from chromatin in DNA-PKcs KO cells. Depletion of FBXL12 in U2OS PKcs-KO resulted in an increase of the intensity of Ku foci induced by ionizing radiation in PKcs-KO cells similarly to what was observed with NEDi (**Fig.3B**, see **Fig.S5A** for the siRNA-mediated depletion). This effect was observed both in and outside of S-phase, however Ku foci were consistently less intense in S-phase, in agreement with the ATM-dependent mechanism operating in this phase of the cell cycle. Adding NEDi to FBXL12 depleted cells had no additional effect on Ku foci intensity, supporting the model that FBXL12 mediates the neddylation-dependent Ku release. The effect of FBXL12 depletion on Ku foci intensity was not observed in DNA-PKcs proficient cells (**Fig.S5B**), indicating that FBXL12 removes Ku only once it accumulates in chromatin. Considering that excessive Ku accumulation in chromatin can be deleterious to other DNA transactions such as transcription or replication, we analyzed the impact of FBLX12 depletion on the fitness of wild-type or DNA-PKcs deficient cells. While depletion of FBXL12 had no measurable effect on wild-type cells, it significantly reduced the cell fitness of DNA-PKcs-deficient cells (**Fig.S5C**), suggesting that excessive Ku accumulation in chromatin is deleterious to cells. This strong effect of FBXL12 depletion on cell fitness was not observed in DNA ligase IV knock-out (**Fig.S5C**) or upon inhibition of DNA-PK kinase activity (**Fig.S5D**). Together, these data identify FBXL12 as the mediator of the neddylation-dependent mechanism preventing Ku accumulation in chromatin in cells deficient for DNA-PKcs.

### CtIP-dependent DNA end resection overcomes Ku entry into chromatin in S-phase

ATM has a dual role in limiting Ku accumulation at single-ended DSBs induced by camptothecin (CPT): (1) it promotes DNA-PKcs removal from ends through phosphorylation of the ABCDE cluster (Britton et al., 2020) and (2) it promotes the initiation of DNA end resection through phosphorylation of CtIP (Chanut et al., 2016). To test whether the role of ATM in antagonizing Ku loading on IR-induced DSBs in S-phase was dependent on DNA end resection, we used siRNA to deplete CtIP, a key activator of DNA end resection (**Fig.3C, see Fig.S5E** for the depletion). Depletion of CtIP in PKcs-KO cells resulted in a strong increase in IR-induced Ku foci intensity in replicating cells treated with NEDi, thereby mimicking the effect of ATM inhibition. Strikingly, inhibiting ATM in the siCtIP conditions did not further increase Ku foci intensity supporting that ATM functions in the same pathway than CtIP to counteract Ku accumulation at DSBs in S-phase (**Fig.3C**). These results suggest that, in S-phase, the ATM-CtIP axis antagonizes Ku overloading induced by the absence of DNA-PKcs, thereby limiting Ku entry into chromatin independently of its neddylation-dependent eviction. In response to IR, DNA resection occurs mainly in the G2 phase of the cycle ((Beucher et al., 2009); **Fig.S5F**, WT+IR condition). However, we observed that both Lig4 or DNA-PKcs loss resulted in IR-induced DSBs being resected in S-phase (**Fig.S5F**), supporting the model that blocking fast NHEJ-dependent DSB repair stimulates DSB end resection.

We also analyzed whether DNA-PKcs prevents Ku overloading on seDSBs induced by CPT in S-phase (**Fig.S5G**). To that aim, WT, PKcs-KO, LIG4-KO and LIG4/PKcs-KO U2OS were treated with CPT in presence of ATMi, to block the CtIP/ATM-dependent mechanisms removing Ku from seDSBs in S-phase. Under these conditions, we confirmed that loss of DNA-PKcs increased Ku foci intensity, by ∼4-times (**Fig.S5G**). Ku overloading did not result from NHEJ inhibition, since it was not observed in response to DNA ligase IV knock-out alone. Altogether, these data support the model that DNA-PKcs functions to block Ku overloading at DSBs throughout the cell cycle, but that, in S-phase, DNA resection is able to quickly overcome Ku overloading when DNA-PKcs is absent.

### DNA-PKcs physically protects transcription at the vicinity of DNA ends

Prior biochemical experiments in cell extracts suggested that excess Ku loading could impede transcription from a nearby promoter *in vitro* (Frit et al., 2000; Ono et al., 1996). To test whether unscheduled Ku entry on DNA can affect transcription at the vicinity of DNA ends in cells, we developed an assay in which U2OS cells were transfected with a linear double-stranded substrate carrying the small Simian Virus 40 promoter (SV40 Pro, 314 bp) followed by the GFP cDNA (720 bp) and the Bovine Growth Hormone poly-adenylation sequence (BGH PolyA, 234 bp, **Fig.4A**). Ku entry onto the promoter or the GFP coding unit is expected to reduce GFP expression. This substrate was co-transfected with a circular mCherry-coding plasmid to restrict the analysis to transfected cells. In the absence of DNA-PKcs, the amount of GFP positive cells was reduced by 50% as compared to WT or to LIG4-KO cells (**Fig.4B**, left panel), demonstrating that the decrease in GFP expression was not due to loss of NHEJ in PKcs-KO cells.

**Figure 4:**
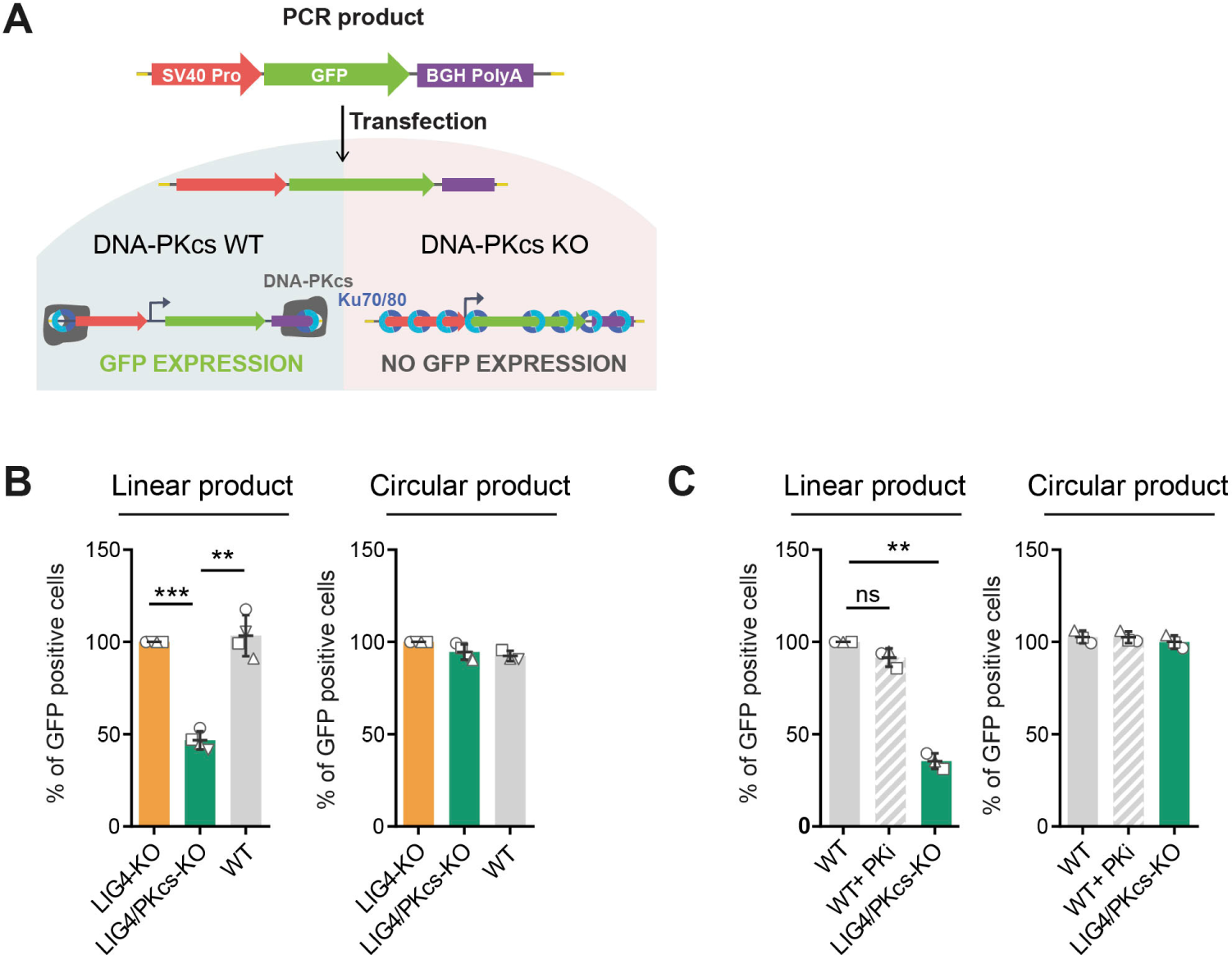
DNA-PKcs deficiency impacts on transcription at the DNA end vicinity. **A.** Scheme depicting the linear substrate used to monitor Ku interference with transcription. Cells are co-transfected with a circular plasmid coding for mCherry and with a linear PCR product with a minimal transcription unit coding for GFP with 5’DNA ends protected against exonuclease by five phosphorothioate linkages or with the same transcription unit inserted in a circular plasmid. Upon transfection, Ku is expected to bind to the linear substrate DNA ends and thread in, physically impeding its transcription. **B.** WT, LIG4-KO or LIG4/PKcs-KO U2OS cells were co-transfected with a mCherry-coding circular plasmid together with the circular (left panel) or linear (right panel) GFP-coding substrates for 24 h before being analyzed by flow cytometry. The graphs represent the percentage of GFP positive cells among the cells successfully transfected, as identified using the mCherry fluorescence. **C.** WT or LIG4/PKcs-KO U2OS cells, treated with 250 nM nedisertib (PKi) when indicated, were co-transfected with a mCherry-coding circular plasmid together with the circular or linear GFP-coding substrates for 24 h before being analyzed by flow cytometry. The graphs represent the percentage of GFP positive cells among the cells successfully transfected, as identified using the mCherry fluorescence.

Importantly, the expression of the same substrate integrated in a circular plasmid was not affected (**Fig.4B**, right panel), showing that DNA ends are required to observe the effect of DNA-PKcs on transcription. In addition, DNA-PK kinase inhibition did not impact the percentage of GFP-positive cells, both with the linear and plasmid substrates (**Fig.4C)**, showing that this function of DNA-PKcs is not mediated by its catalytic activity nor its role in NHEJ. Together these data support that DNA-PKcs presence, but not catalytic activity, protects transcription at the DNA end vicinity.

## Discussion

This study establishes in human cells and in *Xenopus laevis* egg extracts that DNA-PKcs acts as a key regulator of Ku entry into chromatin from DNA ends. This corresponds to a direct, structural role, independent from its kinase activity, and likely relies on DNA-PKcs retaining Ku at the DNA end, occluding further Ku loading (**Fig.5A**). We show this function of DNA-PKcs in restricting Ku loading in two human cell lines using several complementary techniques: by quantifying the fluorescence intensity of individual Ku foci, by analyzing the number and organization of Ku localizations in each focus using STORM and by monitoring Ku association on chromatin using a cell fractionation assay. In addition, we take advantage of a quantitative single molecule assay using *Xenopus* egg extracts to validate and extend our findings that DNA-PKcs enforces a 1:1 Ku-DNA end stoichiometry, and that DNA-PKcs depletion allows additional Ku molecules to thread onto DNA. We also exploited single-molecule analysis to biochemically reconstitute this regulatory function of DNA-PKcs. These analyses support a direct and physical role for DNA-PKcs in blocking additional Ku loading onto DNA.

**Figure 5:**
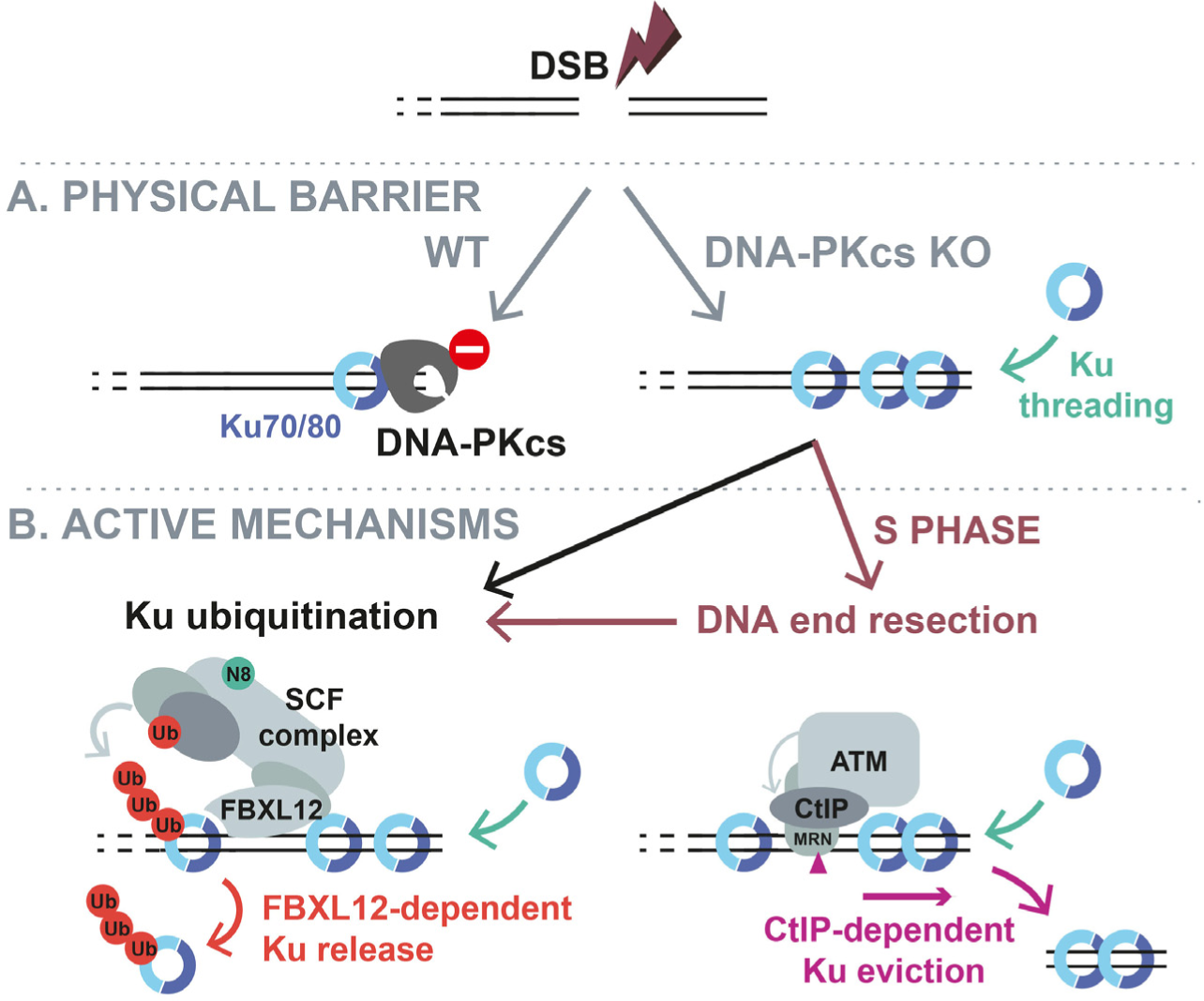
Model summarizing the different barriers to Ku overloading. Upon DSB formation, Ku quickly load on DNA ends. The formation of a Ku-DNA-PKcs complex physically restrains Ku entry into chromatin, enforcing the 1:1 Ku-DNA end stoichiometry (**A)**. In absence of DNA-PKcs, multiple Ku proteins can load and slide from the DNA end into chromatin. The progressive accumulation of larger amounts of Ku into chromatin is actively restricted by two mechanisms (**B**). In all cell cycle phases (bottom left), Ku is evicted *via* its ubiquitination by a FBXL12-containing SCF complex whose activity relies on its neddylation. In S-phase (bottom right), an ATM/CtIP-dependent mechanisms overcomes Ku accumulation through DNA end resection.

This study also establishes that a FBLX12/neddylation-dependent process actively counteracts Ku overloading onto chromatin throughout all phases of the cell cycle, most likely through direct ubiquitination of Ku by a Skp1-Cul1-F-box (SCF) ubiquitin-ligase complex in which FBXL12 is the F-box protein (**Fig.5B**, left panel). The ubiquitination-dependent removal of Ku in this context is supported by the appearance of an ubiquitinated form of Ku both in response to DNA damage in human cells extracts and upon incubation of *Xenopus* egg extracts with DNA ends. In both cases, Ku ubiquitination was blocked by a neddylation inhibitor. What controls FBXL12 recruitment and which signal triggers Ku ubiquitination remain to be investigated. Other E3 ubiquitin ligases have been shown to mediate Ku ubiquitination and/or release, including RNF8, RNF138 and RNF126 (Ishida et al., 2017; Ismail et al., 2015; Kolas et al., 2007) and their contribution in antagonizing Ku accumulation on chromatin remains to be tested.

This study also establishes that ATM/CtIP-dependent resection can overcome Ku accumulation on chromatin in DNA-PKcs deficient cells (**Fig.5B**, right panel). Since most DSBs induced by IR are repaired by NHEJ, this suggests that excessive Ku accumulation resulting from DNA-PKcs loss still allows end resection in S phase. However, as a consequence of Ku molecules covering a larger region of DNA in absence of DNA-PKcs, one might expect that resection is initiated further from the end, since protein blocks play a role in defining the position of the MRX/MRN-dependent incision (Gnugge et al., 2023; Reginato et al., 2017). Supporting this model, loss of DNA-PKcs was found to stimulate the resection of AsiSI-induced DSBs, resulting in larger regions of ssDNA being produced in HCT116 PKcs-KO as compared to WT cells (Zhou et al., 2014). Intriguingly, in the G_0_ phase of the cell cycle, DNA-PK kinase activity is essential for the resection of some DSBs, potentially through phosphorylating CtIP (Deshpande et al., 2020), and in that context depletion of FBXL12 leads to hyper-resection (Fowler et al., 2022). This further supports our finding which shows that DNA end resection can overcome Ku accumulation in chromatin, but potentially at the expense of extended DNA resection.

In addition, our data demonstrate that the absence of DNA-PKcs also leads to Ku overloading at single-ended DSBs induced by CPT when ATM is inhibited. ATM promotes Ku release from seDSBs by stimulating nuclease activities of the CtIP/MRN complex (Chanut et al., 2016) and through direct phosphorylation of DNA-PKcs (Britton et al., 2020). Under these conditions, it was previously shown that seDSBs are repaired unfaithfully by NHEJ, resulting in chromosomal aberrations and cell death, which can be prevented by inhibiting NHEJ (Balmus et al., 2019; Britton et al., 2020). It was postulated that under these conditions (CPT+ATMi+NHEJi), HR still operates, albeit at a slower rate (Balmus et al., 2019). We speculate that excessive Ku accumulation in the absence of DNA-PKcs impacts the ability of HR to proceed. This could explain why the sensitivity to CPT of ATM-deficient mouse embryonic stem cells could be rescued by knocking-out LIG4, XRCC4, XLF or Ku, but strikingly not by inactivating DNA-PKcs (Balmus et al., 2019).

Our findings also raise the question of the length of DNA covered by Ku in the absence of DNA-PKcs and how chromatin boundaries limit Ku spreading or whether they are affected by excessive Ku accumulation. Consistent with chromatin acting as a barrier, inhibition or depletion of KDAC1/2, which promotes chromatin opening, increases Ku loading onto DNA breaks while reducing NHEJ and HR efficiency (Miller et al., 2010). Chromatin boundaries also determine how DNA damage signaling spreads onto chromatin and could therefore impact the ability of Ku to thread onto DNA (Caron et al., 2012). Conversely, Ku threading could impact chromatin organization. In agreement, it was previously shown *in vitro* that Ku can evict the H1 linker histone and “peel off” the DNA from the nucleosome to associate to chromatinized DNA substrates, however this accommodated only 1-2 Ku molecules (Roberts and Ramsden, 2007).

The spreading of Ku molecules into chromatin also impacts the genomic processes occurring in the vicinity of the DNA end. In line with previous *in vitro* studies (Frit et al., 2000; Ono et al., 1996), we established here that the presence of DNA-PKcs is important for enabling transcription of a linear substrate. Since this effect was not dependent on DNA-PK kinase activity and only observed on a linear substrate, without impacting transcription on the same substrate integrated into a circular plasmid, these data support the idea that Ku sliding from the DNA ends in absence of DNA-PKcs impedes transcription. The deleterious impact of Ku accumulation on chromatin is also supported by the specific reduction of cell fitness induced by FBXL12 depletion in DNA-PKcs deficient cells. The lack of DNA-PKcs homolog in several organisms, such as yeast, raises the question of how this protective role of DNA-PKcs is fulfilled there, especially considering that Ku overloading would likely be even more detrimental to the compact yeast genome. It is possible that in yeast the ability of Ku to thread onto DNA is more restricted as suggested in an elegant study in which mutants of yKu70 with an increased ability to slide into DNA were identified (Balestrini et al., 2013). Another possibility is that another Ku partner in yeast substitutes for DNA-PKcs in this function.

To conclude, our work unveils a new and conserved function for DNA-PKcs in regulating the loading of Ku at DNA double strand breaks, along with two active mechanisms which can limit or overcome Ku overloading. Given this role, targeting DNA-PKcs for degradation by small molecules, such as anti-DNA-PKcs PROTACs, or blocking its interaction with Ku could be exploited to induce synthetic lethal situations in cells with specific defects, for example in the FBXL12/neddylation-dependent mechanism that actively extracts Ku from chromatin.

## Supporting information

Supplementary figures with captions

## Acknowledgments

We acknowledge the imaging facility TRI, member of the national infrastructure France-BioImaging infrastructure supported by the French National Research Agency (ANR-10-INBS-04) and Hannah Careless for performing the first analyses with the GFP reporters. This work was supported by the Ligue Contre le Cancer as Equipe Labellisée 2018 (S.B., P. Calsou, P.F.), La Ligue contre le Cancer comité Midi-Pyrénées (to S.B.), la Fondation pour la Recherche Médicale (to M.B.) and the National Institutes of Health grant R01GM115487 (to J.J.L.). We thank Sylvain Cantaloube (CBI, Toulouse) for technical assistance with STORM imaging. P. Calsou is a scientist from INSERM. We thank Johannes Walter and his laboratory for access to their frog facility, for sharing reagents, and for helpful discussions.

## Author contributions

Investigation—generation of human cell models, immnoblotting, imaging, Ku foci quantification, flow cytometry, analysis, funding acquisition, and writing, M.B.; investigation— all experiments using *Xenopus* eggs extracts, analysis, and writing, A.M.; investigation— STORM acquisition and analysis, and writing, A.P.; investigation—flow cytometry with the GFP reporters, analysis, and writing, M.-J.P.; investigation—generation of human cell lines, P.F.; investigation—Ku foci imaging, P.Ch.; investigation—immunoprecipitation from human cell extracts, chromatin fractionations from human cells, analysis, conceptualization, supervision, funding acquisition, and writing, P.Ca.; analysis, conceptualization, supervision, funding acquisition, and writing, J.J.L.; investigation—generated plasmids for human cell line investigations and flow cytometry analysis, conceptualization, supervision, funding acquisition, and writing, S.B..

## MATERIAL AND METHODS

### Cell Culture and Treatments

U2OS and HeLa cells were grown in Dulbecco’s Modified Eagle Medium (Gibco) supplemented with 10 % feal bovine serum (Eurobio), 100 U/ml penicillin (Gibco) and 100 µg/ml streptomycin (Gibco) in a humidified incubator at 37 °C and 5 % CO_2_. For immunofluorescence studies, cells were exposed to the indicated dose of X-Ray irradiation using a calibrated irradiation system Faxitron RX-650 (130 kV, 5 mA). Nedisertib (M3814, DNA-PKi, 0.25 µM, Selleckchem), KU-55933 (ATMi, 10 µM, Selleckchem), MLN-4924 (NEDi, 3 µM, Selleckchem), or NU-7441 (3 µM, Tocris) were added when specified 1 h prior to irradiation. Camptothecin (CPT, Sigma-Aldrich) was used at 1 µM for 1h

### Cell Constructions and Transfections

U2OS PKcs-KO complemented with DNA-PKcs or an empty plasmid were previously described (Britton et al., 2020). U2OS WT, PKcs-KO, KO-LIG4 and KO-LIG4/PKcs were generated using the eSpCas9 (1.1) system with reduced off-target editing (Slaymaker et al., 2016). Cells were transfected with the pCAG-eSpCas9-2A-GFP (generous gift from Jizhong Zou, Addgene plasmid #79145; http://n2t.net/addgene:79145; RRID:Addgene_79145) plasmid co-expressing the S. pyogenes Cas9 variant K848A K1003A R1060A and a guide against LIG4 (target sequence GGGGTAAGAGAACCTTCAGT) and/or DNA-PKcs (target sequence GGTACCCACCCAGCACCGCG) cloned using BbsI digestion. For RNA interference experiments, cells were transfected twice (24 h and 48 h after seeding) with lipofectamine RNAiMAX Reagent (Thermo Fisher Scientific) at a final concentration of 50 nM per well. Experiments were done 48 h after the second round of transfection. Target sequences of siRNA used were: CGUACGCGGAAUACUUCGA for control, UGAUCGAUAUGCUCAUAAA for FBXL12 and GCUAAAACAGGAACGAAUC for CtIP.

### Cell fitness assay

For cell fitness assays, U2OS WT, PKcs-KO or LIG4-KO cells were seeded the day after the 2^nd^ siRNA transfection in 96-well plates. For each cell line, serial two-fold dilutions were performed to seed from 20000 to 156 cells per well. Cells were allowed to grow for 3 days, fixed 1 h at 4°C by adding 10% trichloroacetic acid to reach a 3.33% final concentration and washed with tap water. After drying the plates, cells were stained 30 min at room temperature with 0.057% sulforhodamin B in 1% acetic acid, washed four times with 1% acid acetic and dried again. Finally, the dye was resuspended by a 1 h incubation in 200 µL of 10 mM Tris-base solution, and SRB levels were measured by absorbance at 490 nm using a μQuant microplate spectrophotometer (Bio-Tek Instruments). Percentages of cell growth are expressed after normalization relative to the well with the maximum cell concentration of each cell line transfected with Ctrl siRNA.

### Ku Foci Immunodetection

#### Staining procedure

Immunodetection of Ku foci was performed as previously described (Britton et al., 2013). U2OS and HeLa cells were seeded in 24-well plate on #1.5 glass coverslips (VWR), irradiated and/or treated 24 h later. For STORM experiment, U2OS cells were seeded in 12-well plate on #1.5H glass coverslips (18 mm diameter, Marienfeld, ref. 0117580). Cells were washed twice with PBS and incubated 3 min at room temperature in CSK buffer (10 mM PIPES pH 7.0, 100 mM NaCl, 300 mM sucrose, 3 mM MgCl_2_) containing 0.7% Triton X-100 and 0.3 mg/mL Ribonuclease A (CSK+R). After three PBS washes, cells were again incubated 3 min in CSK+R. Cells were then washed three times and fixed with 2% paraformaldehyde in PBS for 20 min at room temperature. Cells were washed with PBS, permeabilized 5 min with PBS containing 0.2% Triton X-100, washed again and incubated 10 min at room temperature in blocking buffer (PBS 0.1% Tween-20, 5% bovine serum albumin). Cells were incubated 75 min at room temperature with primary anti-Ku and/or PCNA antibody diluted in blocking buffer, washed with PBS 0.1% Tween-20 (PBS-T), and incubated again 45 min with secondary antibody coupled to respectively Alexa Fluor 488 or 594 in blocking buffer. The list of primary and secondary antibodies used here with their conditions of use is provided in **Table 2**. Cells were finally washed with PBS 0.1% Tween-20 and stained with 0.2 µg/mL DAPI in PBS for 15 min at room temperature. Coverslips were mounted with Vectashield (Vector laboratories). For STORM experiments, AlexaFluor 647-coupled goat anti-mouse antibodies were used to immunodetect Ku.

#### 3D-SIM imaging and analysis

Images were acquired with a Zeiss Elyra 7 3D Lattice SIM super-resolution microscope with a 63x objective (PLANAPO NA 1.4, Zeiss) and dual sCMOS cameras (pco.edge). 3D-SIM reconstructions were performed on 2 µm depth-images (interval 0.091 µm) with Zen Blue 3.3 (Zeiss). Threshold for PCNA positive/negative cell was chosen each experiment based on the cell population geometric mean of nuclear PCNA-integrated density staining. Ku foci number and intensity quantifications were performed on maximum intensity projection using the find maxima function of Image J software (v1.53t). To circumvent quantification bias associated with large intensity variation, each nucleus was automatically contrasted based on the image’s histogram (auto B&C function) and the returning maximal threshold value was used to adjust the prominence of the find maxima function. The prominence was calculated using the following experimentally determined two-phase decay model: *f*(*x*) = 1132 + ((18.66 - 1132) x 8.908 x 0.01) x *exp*(-0.00252*x*) + ((18.66 -1132) x (100 - 8.908) x 0.01) x *exp*(-0.00005977*x*).

#### STORM imaging and analysis

3D Stochastic Optical Microscopy (STORM) was performed on an inverted i-SPT Nikon Ti-E/B microscope equipped with an Apo TIRF 100X, NA1.49 oil DIC objective and an adaptive optics system (MicAO 3D-SR, Imagine Optic) that was used to introduce an optical astigmatism during image acquisition. Excitation was performed in HILO configuration with a 647 nm laser line (300 mW) from MPB Communications Inc. The excitation light signal was filtered by a quad-band dichroic mirror (Nikon N-STORM TIRF Filter Set, 97335) whereas the emission light was filtered by 705/72 nm filter (Chroma, ET705/72 nm) and detected on a iXon Ultra DU897 EM-CCD camera (Andor). Microscopy slides were fitted with a 13 mm x 0.6 mm Coverwell modular hybridization system (Citi-Fluor EMS) that was filled with STORM buffer (Smart Buffer Kit, Abbelight) and sealed with a glass coverslip coated with TetraSpeck™ 0.1 µm beads (ThermoFischer, T7279). Image acquisition was performed for 30,000 frames at 512 x 512 pixels with an exposure time of 30 ms at full power of the 647 nm laser. After acquisition, the localization of individual fluorophores was performed with the ThunderSTORM plugin (Ovesny et al., 2014) on ImageJ (Schneider et al., 2012). Individual localizations with an uncertainty above 40 nm and intensity above 5000 were discarded from further analysis. The remaining localizations were loaded into the Point Cloud Analyst software (Levet and Sibarita, 2023) for visualization and cluster analysis purposes. For the identification of individual clusters we used the SR-Tesseler method of the PoCA software with the following settings: minimal number of localizations = 10, maximal number of localizations = 10000, Cut distance = 100 nm, coefficient factor = 2. The characteristics of each cluster (size, volume and number of localizations) were exported and used for further analysis in GraphPad Prism. For visualization of individual clusters, only the localizations within identified clusters were used for rendering using the Heatmap function of the PoCA software.

### Whole Cell Extracts

Cells were washed with PBS, scrapped in a lysis buffer containing 120 mM Tris-HCL pH 6.8, 4% SDS, 20% glycerol and further lysed with 10 strokes through a 24G needle. Protein concentration was determined with a Nanodrop™ by measuring absorbance at 280 nm. Immunoblotting, as described below, was then performed.

### Cell Fractionation Assay

Stock solution of Calicheamicin γ1 (Cali), gift from P. R. Hamann (Wyeth Research, Pearl River, NY, USA), was made at 40 µM in ethanol and stored at -20°C. For drug-exposure, exponentially growing cells in 60 cm diameter dishes were either mock-treated or treated with Cali in fresh medium for 1 hr at 37 °C. Then cells were washed with phosphate-buffered saline (PBS) and trypsinized. Pellets were fractionated as follows. Cells were first resuspended for 7 min on ice in 120 µL of extraction buffer 1 (50 mM Hepes pH 7.5, 150 mM NaCl, 1 mM EDTA, 0.1% Triton X-100, 1X Halt protease and phosphatase inhibitor cocktail (Thermo Fisher Scientific)) with intermittent gentle vortexing. Following centrifugation at 14 000 rpm for 3 min, the supernatant 1 was removed and stored, pellets were gently resuspended with pipette tips in 120 µL of extraction buffer 2 (50 mM Hepes pH 7.5, 75 mM NaCl, 1 mM EDTA, 0.025% Triton X-100, U RNAseA/T1 (Thermo Fisher Scientific)), incubated for 15 min at 25°C under agitation and centrifuged as above. Supernatants 2 were pooled with supernatants 1 (soluble protein fraction). Pellets (chromatin fraction) were resuspended in 120 µL lysis buffer (50 mM Tris pH 8.1, 10 mM EDTA, 0.1% Triton X-100, 0.3% SDS) and sonicated (Vibracel, Bioblock Scientific). Protein content was measured with BCA reagent (Pierce). Immunoblotting was then performed as described below.

### Ku Immunoprecipitation from Human Cells

M280 anti-mouse magnetic beads were coupled to control mouse IgG2a (Dako) or mouse monoclonal anti-Ku antibodies (Thermo Scientific, clone 162) with a ratio of 1.25 µg protein to 50 µL beads suspension, in PBS 0.1% Tween-20 (Sigma-Aldrich) (PBS-T) under rotation at 4°C overnight, followed by 3 washes in PBS-T and storage at 4°C in PBS-T supplemented with sodium azide 0.02%. For immunoprecipitation, 10 µL of anti-Ku or control magnetic beads suspension were washed 3 times in PBS-T, dried over a magnet and incubated in 250 µL microtubes (Eppendorf) under rotation at 4°C overnight in 100 µL extraction buffer 1 with 30 µg of proteins from soluble fraction as indicated, then transferred to clean tubes, washed 3 times in PBS-T, resuspended in lysis buffer supplemented with 1X loading buffer (50 mM Tris– HCl pH 6.8, 10% glycerol, 1% SDS, 300 mM 2-mercaptoethanol, 0.01% bromophenol blue) and denaturated 5 min at 95°C before protein separation by SDS–PAGE on 4-15% precast gels (Biorad).

### Immunoblotting

Equivalent protein amounts were denaturated in loading buffer at 1X final concentration at 95°C for 5 min, separated on SDS-PAGE gels (BioRad 4-15% TGX pre-cast gels) before transfer overnight onto Immobilon-P polyvinylidene difluoride (PVDF, Millipore) or nitrocellulose (0.45 µm pore, Bio-Rad or Protran, GE Healthcare) membranes. Staining with AdvanStain Iris (Advansta), or Ponceau S, controlled homogeneous loading and prestained protein ladder allowed cutting the membrane into stripes to simultaneously blot multiple antibodies. Membranes were blocked for 60 min with 5% non-fat dry milk in PBS-T buffer, incubated as necessary with primary antibody diluted in PBS-T containing 1% or 2.5% bovine serum albumin (immunoglobulin- and lipid-free fraction V, Sigma-Aldrich) and washed 3 times with PBS-T; membranes were incubated for at most 1 h with HRP-conjugated secondary antibodies in PBS-T and washed three times with PBS-T. A list of antibodies and conditions of use is provided in **Table 2**. Immuno-blots were either visualized using autoradiography films together with enhanced chemiluminescence (WesternBright ECl, Advansta), or by imaging membranes on an Amersham Imager 600 (GE Healthcare) with HyGLO Quick Spray (Denville Scientific).

### GFP Reporters Preparation and Flow Cytometry Analysis

The pUC18-GFP plasmid (will be deposited on Addgene) was produced by cloning using a EcoRI+BamHI digestion a minimal transcription unit for GFP expression, consisting of the small SV40 promoter, the GFP ORF and the BGH polyadenylation signal, generated by PCR assembly. For use as a control, a pUC18-LucFF plasmid (will be deposited on Addgene) was also generated by replacing the GFP ORF by the firefly luciferase ORF. The linear GFP substrate was generated from the pUC18-GFP by PCR with DreamTaq (Thermo Fisher Scientific) according to manufacturer’s instructions using the following amplification program: 95°C 30s, 36 cycles (95°C 30s, 68°C 30s, 72°C 30s) and 72°C 5 min and the phosphorothioate-modified SV40-F 5’-G*G*C*G*A*ATTCCTGTGGAATGTGTGTCAGTTAGGGTGTGG-3’ and BGH-R 5’-C*G*C*G*G*ATCCCCATAGAGCCCACCGCATCCC-3’ primers (* indicates phosphorothioate linkages, provided by Eurofins). The PCR product was purified after migration on a 1% agarose TAE 0.5X gel using the Wizard PCR Clean-Up system (Promega) according to manufacturer’s instructions. For transfections, U2OS cells were seeded in 60 mm dishes (800,000 cells for U2OS WT and LIG4-KO, and 900,000 cells for LIG4/PKcs-KO) 24 h prior transfection using lipofectamine 2000 (Thermo Fisher Scientific) following manufacturer’s instructions. 4.9 μg DNA was transfected per dish containing a 1:1 molar ratio of pmCherry-N1 (Clontech) and of the pUC18-GFP or linear GFP substrate. 24h after transfection, cells were trypsinised, collected in cold medium and then centrifuged at 4°C at 1000 RPM for 5 min. Cells were resuspended in cold PBS containing 1% bovine serum albumin (PBS-1% BSA), centrifuged at 1000 RPM at 4°C for 5 min and fixed by incubation for 15 min at room temperature in 2% paraformaldehyde in PBS. After addition of PBS-1% BSA, the cells were centrifuged at 1000 RPM at 4°C for 5 mins, washed in PBS-1% BSA and resuspended in PBS 1% BSA before analysis on LSRII Fortessa X20 (Becton Dickinson). A minimum of 300,000 cells were acquired per condition. Data were analyzed and formatted using FlowJo v10.8.1.

### Flow cytometry

1.10^6^ cells were seeded in 60 mm dishes and treated 24h later as specified. Cells were washed with PBS, trypsinised, collected with cold DMEM, and centrifuged 5 min with 400g at 4°C. Cells were again washed with cold PBS, centrifuged 4 min with 400g at 4°C and pre-extracted on ice for 10 min with PBS containing 0.2% Triton TX-100. After PBS-1% BSA addition, cells were fixed 15 min at RT in 500 µL PBS containing 2% paraformaldehyde. PBS-1% BSA was added to stop the reaction, cells were washed with PBS-1% BSA and resuspended in 100 µL of PBS-T 5% BSA containing primary antibody (see **Table 2** for antibodies and dilution) for 1h at RT. After PBS-1% BSA addition, cells were incubated 30 min in 200 µL of PBS-T 5% BSA containing secondary antibody (see **Table 2** for antibodies and dilution) at RT and obscurity. PBS 1%-BSA was added in each condition and cells were finally incubated at least 30 min in PBS containing 0.25 mg/mL RNAse A and 2 µg/ml DAPI. A minimum of 40,000 cells were processed by a BD LSRII flow cytometer (Becton Dickinson). FlowJo v10.8.1 was used to analyze and format data.

### Egg Extract Preparation and Immunodepletion

High-speed supernatant (HSS) of egg cytosol was prepared as described previously (Lebofsky et al., 2009). The female frogs used to produce oocytes were cared for by the Center for Animal Resources and Comparative Medicine at Harvard Medical School (AAALAC accredited). Work performed for this study was in accordance with the rules and regulations set by AAALAC. The Institutional Animal Care and Use Committee (IACUC) of Harvard Medical School approved the work.

For DNA-PKcs immunodepletion, antibodies, described below, were bound to protein A sepharose beads at a ratio of 20 µg antibody per 1 μL beads. For Ku80, antibodies were bound to the beads at a ratio of 4 µg antibody per 1 μL beads. For XLF and XRCC4 depletions, antibodies were bound to the beads at a ratio of 3 µg antibody per 1 μL beads. Beads were extensively washed with egg lysis buffer with sucrose (ELBS; 10 mM HEPES, pH 7.7, 50 mM KCl, 2.5 mM MgCl2, 250 mM sucrose). Extract was supplemented with nocodazole to a final concentration of 7.5 ng/ml and one volume of extract was incubated for 3 rounds with 1/3 volume (for DNA-PKcs) or 1/5 volume (for Ku80) of antibody-bound beads for 60 min per round on a rotator at 4°C. Beads were pelleted by centrifugation at 2000 g after each round, and extract was collected with a standard P200 pipette tip followed by an ultrafine gel-loading tip. The immunodepleted extract was then centrifuged for 5 minutes 16000 r.f.c. at 4°C and supernatant was transferred to a new tube.

### Halo-Ku80/Ku70 Preparation

#### Plasmid Generation

N-terminally Halo-tagged Ku80 was cloned into pFastBac1 by PCR amplifying Halo tag coding sequence from pTG344 using primers oAM082 and oAM083 and PCR amplifying pFastBac1 plasmid pTG246 containing *X. laevis* Ku80 (Graham et al., 2016) using primers oAM084 and oAM085, followed by Gibson assembly with HiFi DNA Assembly Master Mix (Invitrogen). To generate N-terminally His-Halo tagged Ku80 in pACEBac1, Halo-tagged Ku80 was PCR amplified from pAM047 using primers oAM107 and oAM108 and inserted into pACEBac1 cut with NotI by Gibson assembly. The cDNA encoding *X. laevis* Ku70 was cloned into pIDK by PCR amplifying Ku70 from pTG284 using primers oAM111 and oAM112 and inserted into pIDK cut with XhoI and KpnI by Gibson assembly. The His-Halo-Ku80 DNA and Ku70 were cloned into a single expression plasmid (pAM086) using the MultiBac system (Trowitzsch et al., 2010). The bacmid encoding Halo-Ku80/Ku70 complex was obtained by electroporating pAM086 into DH10EMBacY electro-competent cells and purified using ZR BAC DNA miniprep kit (Zymo Research).

#### Purification of His-Halo-Ku80:Ku70

Baculovirus encoding Halo-Ku80/Ku70 was amplified in three stages (P1, P2, and P3) in Sf9 cells (Expression Systems). To express *X. laevis* Halo-Ku80/Ku70, Sf9 cells at a density of 2 x 10^6^/mL in 500 mL of Sf900 III serum-free medium (Invitrogen) were infected with 5-10 mL P3 baculovirus (MOI > 1). Cells harvested 60 hours post-infection were pelleted at 500 x g for 15 min, washed with phosphate-buffered saline (PBS; 135 mM NaCl, 2.7 mM KCl, 4.3 mM Na_2_HPO4, 1.4 mM KH_2_PO4; Teknova), and centrifuged again for 5 min at 500 g at 4°C. Pellets were flash-frozen in liquid nitrogen and stored at -80°C. Thawed cell pellets were resuspended in ∼5 volumes lysis buffer (20 mM Tris-HCl pH 8, 1 M NaCl, 20 mM imidazole, 10 % glycerol, 0.2% Triton X-100, 5 mM β-mercaptoethanol, 1 mM phenylmethylsulfonyl fluoride, and cOmplete protease inhibitor cocktail tablet (Sigma-Aldrich)). Cells were lysed by sonication on ice and the insoluble fraction was pelleted via centrifugation for 1 hr at 40,000 g at 4°C. The clarified lysate was incubated with 0.5 mL NiNTA resin (QIAGEN) for 1 hr at 4°C on a rotary. The resin was washed 3 times with 10 mL of lysis buffer in a disposable column. The protein was eluted in 10 rounds with 500 µL/each of Lysis buffer + 250 mM Imidazole; the elutions were pooled and concentrated using 50kDa MWCO centrifugal concentrator (Millipore). Recombinant Halo-Ku80/Ku70 was further purified on a HiTrap Q HP 1 mL column (Cytiva) connected to an AKTA Pure FPLC with a 100-1000 mM NaCl gradient in 25 mM Hepes pH 7.8, 10% glycerol, 1 mM DTT, 0.05 mM EDTA buffer. The eluted protein was buffer exchanged into storage buffer (25 mM Hepes pH 7.8, 300 mM NaOAc, 1 mM DTT, 0.5 mM EDTA, 10 % glycerol) using 50 kDa MWCO centrifugal concentrator (Millipore), concentrated to ∼10 µM, frozen in liquid nitrogen, and stored at -80°C.

#### Fluorescence labeling of Halo-Ku80/Ku70

A fivefold molar excess of sulfo-Cy5-Halo ligand in DMSO was added to Halo-Ku80/Ku70 protein in its storage buffer. Preparation of sulfo-Cy5-Halo substrate was previously described (Graham et al., 2017). The mixture was incubated for 20 min at 4°C on a rotator and then centrifuged at 16,000 r.c.f. at 4°C. Labeled protein was separated from free dye on a Superdex 200 Increase 10/300 column equilibrated with storage buffer. Peak fractions were pooled and concentrated with a 50 kDa MWCO centrifugal concentrator. To determine protein concentrations absorbance at 280 nm was corrected for Cy5 absorbance at 280 nm and calculated assuming an extinction coefficient of 129,150 M^−1^ cm^−1^ at 280 nm. Dye concentrations were calculated according to absorbance at 650 nm for Cy5 (assuming an extinction coefficient of 250,000 M^−1^ cm^−1^). The degree of Halo-Ku80/Ku70 labeling was obtained by division of the dye molar concentration by the protein molar concentration.

### Purification of endogenous DNA-PKcs from egg extract

A biotinylated 19 bp duplex was prepared by combining 6 nmol each of oAM094: 5’-/5Biosg/CGTTTTTCCATAGGCTCCG and oAM571: 5’-CGGAGCCTATGGAAAAACG (Integrated DNA Technologies), in 200 µL anneal buffer (20 mM Tris-HCl, 300 mM NaCl, 1 mM EDTA), and placed in a 2 L beaker containing water heated to 85°C that was allowed to cool overnight. Streptavidin-sepharose beads (300 µL, Cytiva) were washed twice with 2x Bead Wash Buffer (10 mM Tris, pH 7.4, 2 M NaCl, and 20 mM EDTA), and resuspended in 0.5x Bead Wash Buffer containing 1.32 nmol of biotinylated DNA and rotated for 1 h at 4°C. The DNA-bound magnetic beads were then extensively washed with Bead Wash Buffer, followed by washing with Blocking Buffer (10 mM Hepes, pH 7.7, 50 mM KCl, 2.5 mM MgCl_2_, 250 mM sucrose, and 0.02 % Tween-20). Beads were then resuspended in 500 µL Blocking Buffer. Prepared HSS stored at -80°C was thawed, 4 aliquots of 1 mL, supplemented with nocodazole to a final concentration of 7.5 ng/mL, and centrifuged for 10 min at 16800 rfc in a 4°C chilled centrifuge. The supernatant was transferred to a new tube and centrifuged for an additional 10 minutes at 16800 rfc. The supernatant from the extract was diluted with 3.5 mL of Blocking Buffer, mixed with 0.5 mL of DNA-beads, and placed on ice, inverting the tube every 30 seconds to keep DNA-beads in suspension during a 4 min incubation. The DNA beads were then pelleted by centrifugation at 400 g for 30 s in a chilled centrifuge. The beads were resuspended in 1 mL cold Blocking Buffer and transferred to a polypropylene column. The DNA beads were washed with 10 mL of cold Blocking Buffer at 4°C followed by washing with 8 mL of cold Blocking Buffer containing 100 mM NaOAc at 4°C. DNA-PKcs was eluted from the DNA beads using a gradient of 150 to 600 mM NaOAc in Blocking Buffer. Q-sepharose fast flow resin, 500 µL (GE Healthcare), was washed twice with 2x Bead Wash Buffer (10 mM Tris, pH 7.4, 2 M NaCl, and 20 mM EDTA), followed by extensive washing with Q-dilution Buffer (20 mM Tris, pH 8.0, 10 % sucrose, 0.02 % Tween-20, and 10 mM EDTA). Fractions containing DNA-PKcs were pooled and concentrated to a final volume of 50 µL using 100 kDa MWCO spin concentrator (Millipore) centrifuging 5000 rfc at 4°C. The concentrated eluate from DNA-sepharose resin, was diluted with 5 mL of Q-dilution Buffer, mixed with 500 µL of Q-sepharose resin, and incubated on ice for 4 min. The resin was then transferred to a polypropylene column (Qiagen) and washed with 8 mL Q-binding Buffer (20 mm Tris, pH 8.0, 40 mM NaOAc, 10 % sucrose, 0.02 % Tween-20, and 10 mM EDTA). DNA-PKcs was eluted from the Q resin using a gradient of 100 to 600 mM NaOAc in Q-binding Buffer. Fractions containing DNA-PKcs were pooled and concentrated to a final volume of 50 µL using 100 kDa MWCO spin concentrator (Millipore) centrifuging 5000 rfc at 4°C. Purified DNA-PKcs was stored on ice at 4°C and used within the next 24 h.

### Bulk End-Joining Assay

#### Substrate Preparation

Plasmid pAM089 was linearized by cleavage with EcoRI-HF (New England Biolabs), separated on a 1x TBE agarose gel and extracted by electroelution. DNA was labeled by fill-in of EcoRI overhangs with the DNA polymerase I Klenow fragment (New England Biolabs) in the presence of [α-32P] dATP (Perkin Elmer), dTTP, dCTP, and dGTP incubating for 15 min at 25°C. Labeled DNA was purified using a spin column PCR purification kit (Qiagen) and eluted with 10 mM Tris-HCl, pH 8.0.

#### End-Joining Assay

The bulk end-joining assay was performed as described previously (Graham et al., 2016). Briefly, extracts were supplemented with the following (final concentrations in parentheses): closed-circular DNA pAM089 (30 ng/μL) and ATP regeneration mixture ATP (3 mM); phosphocreatine (15 mM); creatine phosphokinase (0.01 mg/mL; Sigma). For rescue experiments, extract was supplemented with Halo-Ku80/Ku70 (300 nM). For experiments to examine effect of Cullin inhibitor on end-joining extracts were supplemented with either Cullin neddylation inhibitor MLN4924 (200 μM) or DMSO vehicle control and incubated on ice for 5 min. The mixture was incubated for 5 minutes at room temperature and then placed back on ice. To initiate the reaction 1.0 μL of 20 ng/μL radiolabeled linear substrate DNA was added to the extract, and 2 μL sample (0 min) was withdrawn while the reactions were on ice and mixed with 5 µL stop solution (80 mM Tris [pH 8], 8 mM EDTA, 0.13% phosphoric acid, 10% Ficoll, 5% SDS, 0.2% bromophenol blue) and 1 μg proteinase K. Reactions were transferred to room temperature, and additional 2 μL samples were withdrawn at the indicated times and mixed with 5 μL stop solution and 1 μg proteinase K. Samples were digested at 37°C for a minimum of 1 hr, and products were separated by electrophoresis on a 13 Tris-borate-EDTA, 0.8% agarose gel. Gels were sandwiched between filter paper and a HyBond-XL nylon membrane (GE Healthcare), dried on a gel dryer, and exposed to a phosphorscreen, which was imaged with Typhoon FLA 7000 imager (GE Healthcare Life Sciences).

### DNA Pulldown Assay

The DNA pulldown assay was largely performed as previously described (Carney et al., 2020). The protocol with modifications is briefly described below. A Cy3-labeled 100 bp duplex with biotin molecules attached to the 5’ termini, generated for single-molecule experiments, was the DNA substrate used in pulldown experiments. Streptavidin-coated magnetic beads (M-280, Sigma) (96 μL per biological replicate) were washed twice in 2x Bead Wash Buffer (10 mM Tris, pH 7.4, 2 M NaCl, and 20 mM EDTA), and resuspended in 1x Bead Wash Buffer containing 1.1 pmol of biotinylated DNA and rotated for 20 min at 4°C. The DNA-bound magnetic beads were then washed 3x with 2x Bead Wash Buffer and twice with Blocking Buffer (10 mM HEPES, pH 7.7, 50 mM KCl, 2.5 mM MgCl2, 250 mM sucrose, and 0.02% Tween20). Beads were then resuspended in 120 μL of Blocking Buffer and aliquoted, 10 μL per a sample. Extract was supplemented with the ATP regeneration mixture, ATP (3 mM); phosphocreatine (15 mM); creatine phosphokinase (0.01 mg/mL; Sigma-Aldrich), and 30 ng/mL of circular pBlueScript II plasmid to act as carrier DNA (Lebofsky et al., 2009). To maximize ubiquitination signal, extracts were supplemented with p97 inhibitor NMS-873 (200 μM). Effects of Cullin inhibition on Ku80 ubiquitination were examined by supplementing extracts with either Cullin neddylation inhibitor MLN4924 (200 μM) or DMSO vehicle control. The extracts were then incubated on ice for 5 min. To initiate assembly of the NHEJ machinery, the DNA-beads sample was mixed with an equal volume of extract and incubated at room temperature. At a given timepoint the reaction was then layered over 200 μL of Sucrose cushion (10 mM HEPES, pH 7.7, 50 mM KCl, 2.5 mM MgCl2, 500 mM sucrose) in Beckman microfuge tube (5 x 44 mm) and centrifuged for 1 minute at 16800 rfc in a swinging bucket centrifuge (Ependorff, 5430R). The sucrose-cushion was aspirated off and the pelleted DNA-beads were then washed in 200 mL of Egg Lysis Blocking Buffer and resuspended in 20 mL of 1x reducing Laemmli sample buffer. Extract was diluted 1:40 in 1x reducing Laemmli sample buffer to be used as input sample for western blotting. Samples were separated on a 4–15% precast SDS-PAGE gel (BioRad) for 30 min at 200V and western blots were performed as outlined above.

### Single-molecule Stoichiometry Imaging and Analysis

#### Substrate preparation

The Cy3-labeled biotinylated 100 bp duplex was prepared by combining 1 nmol each of oTG048F, oTG415, oTG532, and oTG533 in 50 μL anneal buffer (20 mM Tris-HCl, 300 mM NaCl, 1 mM EDTA) and placed in a 2-liter beaker containing water heated to 90°C that was allowed to cool overnight. Nicks were sealed by adding 10 μL 10x T4 DNA ligase buffer and 2 μL T4 DNA ligase (New England Biolabs) and incubating overnight at 16°C. The ligated product was separated on a 0.5x TBE, 8% polyacrylamide (19:1 bis-acrylamide:acrylamide) gel running at 200 V, 4°C. The band corresponding to the full-length 100 bp DNA was observed by UV shadow and excised from the gel. The gel slice was cut into small fragments suspended in 500 μL 1x TE, and DNA was electroeluted using Elutrap System (Whatman). The DNA eluate was lyophilized and suspended in TE.

#### Preparation of Calibration substrate

The calibration substrate used for channel alignment consists of Cy5 and Cy3 labeled 60 bp duplex and was made as previously described (Fan et al., 2022).

#### Flow Cell Preparation

Coverslips were prepared as previously described (Fan et al., 2022). Briefly, glass coverslips were cleaned by sonication in methanol, washed with DI H2O, and incubated for 1 hour with piranha solution (3:1 mixture of sulfuric acid and 30% hydrogen peroxide). The piranha solution was removed, and coverslips were again washed with DI H2O. The coverslips were then functionalized with a mixture of methoxypolyethylene glycol succinimidyl valerate, MW 5,000 (mPEG-SVA-5000; Laysan Bio, Inc.) and biotin-methoxypolyethylene glycol-succinimidyl valerate, MW 5,000 (biotin-PEG-SVA-5000; Laysan Bio, Inc.) as previously described (Graham et al., 2017).

Microfluidic chambers were assembled as follows: 4.5 mm wide piece of double-sided SecureSeal Adhesive Sheet (Grace Bio-Labs) was placed parallel on each side of holes drilled 10 mm apart in a glass microscope slide. A functionalized coverslip was then secured to the other side of the adhesive sheet and the edges of the coverslip were sealed with epoxy (Devcon). The microfluidic chamber was assembled by inserting PE20 tubing into one hole, PE60 tubing into the other (Intramedic), and fixing in place with epoxy. The microfluidic cells were stored under vacuum until time of use.

#### Single-molecule imaging

A through-objective TIRF microscope configured around an inverted Olympus IX-71 microscope was used to image fluorescent molecules. Laser beams 532 nm (Coherent Sapphire 532) and 641 nm (Cube 641) were expanded and then combined using dichroic mirrors. The colinear beams were expanded and focused onto the rear focal plane of an oil-immersion objective (Olympus UPlanSApo, 100 3; NA, 1.40). For TIRF illumination a focusing lens was translated in the vertical plane. A multipass dichroic mirror separated emission from excitation light, followed by a StopLine 488/532/635 notch filter (Semrock) to further reduce excitation light. A home-built beamsplitter (Graham et al., 2017) separates emission from Cy3 and Cy5 to image on separate halves of an EMCCD camera (Hamamatsu, ImageEM 9100-13) operating at maximum EM gain. The focus was adjusted manually, and the sample was positioned on the microscope using an automated microstage (Mad City Labs).

#### Calibration Data Collection

The flow cell chamber was passivated with 30 µL PBS-BSA buffer (1.0 mg/mL BSA (NEB) in PBS) for 5 min, followed by 5 min incubation with 25 μL 0.2 mg/mL streptavidin in PBS. The flow cell was then washed with PBS-BSA. The calibration substrate, 5-prime biotinylated 60 bp dsDNA labeled with Cy5 and Cy3, was diluted to ∼80 pM in PBS, containing oxygen scavenging system and triplet-state quencher system: protocatechuic acid (PCA; 5 mM; Sigma), protocatechuate 3,4-dioxygenase (PCD; 0.1 mM; Sigma), and Trolox (6-Hydroxy-2,5,7,8-tetramethylchromane-2-carboxylic acid; 1 mM; Sigma). The biotinylated, fluorescent DNA substrate was immobilized on a glass coverslip in a microfluidic chamber. Images were acquired of different fields of view (∼120) with 0.5 sec simultaneous exposure to 532 and 641 nm lasers using a surface power density of 4 mW/cm^2^ for the 532 nm laser and 2.4 mW/cm^2^ for the 641 nm laser.

#### Single-molecule Stoichiometry

The flow cell chamber was passivated with PBS-BSA buffer (1.0 mg/mL BSA; NEB) in PBS for 5 min, followed by 5 min incubation with streptavidin (0.2 mg/ml; Sigma) in PBS. The flow cell was then washed with PBS-BSA and the 5-prime biotinylated 100 bp dsDNA labeled with Cy3 (∼80 pM) in PBS and immobilized on the glass coverslip. The chamber was then washed with egg lysis buffer (ELB wash; 10 mM HEPES, pH 7.7, 50 mM KCl, 2.5 mM MgCl_2_). For extract Halo-Ku80/Ku70 stoichiometry experiments, extracts were supplemented with the following (final concentrations in parentheses): either Cullin neddylation inhibitor MLN4924 (200 μM) or DMSO vehicle control and incubated on ice for 5 min. To Ku-immunodepleted extract, Cy5 labeled HaloKu80/Ku70 (20 nM), closed-circular DNA pAM089 (30 ng/ μL), ATP (3 mM), phosphocreatine (150 mM), and creatine phosphokinase (0.01 mg/ml; Sigma) was added and incubated for 5 min at room temperature. The reaction mixture was then placed on ice and supplemented with oxygen scavenging system and triplet-state quencher PCA (4 mM), PCD (0.08 μM), and Trolox (1 mM). The flow cell was then washed with ELB wash containing PCA (5 mM), PCD (0.1 μM), and Trolox (1 mM). Extract was introduced to the chamber and incubated for 60 min. The flow cell was then washed with ELB wash containing PCA (2.5 mM), PCD (0.05 μM), and Trolox (1 mM). Images were then taken continuously at a rate of 20 frame/s for ∼90 s, alternating between four frames of 641 nm excitation and one frame of 532 nm excitation with surface power densities of 4 mW/cm^2^ for 532 nm and a 641 nm laser power that was adjusted for photobleaching >95 % of Halo-Ku80/Ku70 by the end of the movie. Experimental conditions for *in vitro* minimal reconstitutions involving Halo-Ku80/Ku70 were the same as above except extract were replaced with ELB wash buffer and Cy5 labeled HaloKu80/Ku70 (20 nM) was incubated with 60 nM of NHEJ factors in ELB wash containing 10 % volume storage buffer (20 mM Tris, pH 8.0, 300 mM NaOAc, 10 % sucrose, 0.02 % Tween-20, and 10 mM EDTA) on ice for 5 min before addition to flow cell. Endogenous DNA-PKcs was purified as described above, recombinant XLF and Lig4-XRCC4 were purified as previously described (Carney et al., 2020, Stinson et al., 2022). For each experiment, Halo-Ku80/Ku70 photobleaching was recorded for a minimum of 5 different fields of view.

#### Single-molecule Stoichiometry Image Analysis

Channel alignment of the 532 nm and 641 nm channels was done as previously described (Fan et al., 2022). Briefly, a custom MATLAB script was used to identify fluorescent molecules in the 532 and 641 emission channels based on local maximum and particles were fit using a 2D Gaussian. For each image, spots whose coordinates are greater than 6 pixels from surrounding molecules and are within 6 pixels of a molecule in the corresponding channel are added to the calibration list. The calibration list is then refined based on spot diameter and amplitude. A final calibration list is made using a previously described method (Friedman and Gelles, 2015). Briefly, for each reference image spot coordinates (x_1_,y_1_) in the 532 nm emission channel are mapped onto coordinates (x_2_,y_2_) in 641 emission channel to identify a matching partner spot using the transformation Eq (1):

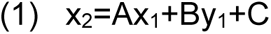

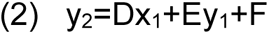

where A–F are fit parameters. Because the field of view is non-uniform, the fit parameters are determined for each spot coordinate (x_1_, y_1_) by fitting pairs of corresponding points from the initial calibration list that are within a defined pixel box from (x_1_, y_1_). A spot that maps greater than 3 pixels from all spots in 641 nm channel is removed from the calibration list. This process is repeated for spots in 641 nm channel, mapping coordinates onto 532 nm channel using the updated calibration list. The 641 nm Channel spots that are within 3 pixels of to its 532 nm channel partner are retained in the calibration list. The calibration list is further refined by iteratively repeating the above process while reducing the pixel threshold from 3 to 0.5 pixels.

#### Detection and Quantification of Halo-Ku80/Ku70 stoichiometry

Analysis of single-molecule Cy5 labeled Halo-Ku80/Ku70 stoichiometry were performed using the following steps. The ROIs for tethered DNAs were identified after averaging the first 5 images with 532 nm excitation in the Cy3 channel using the method described in *Alignment of the Cy3 and Cy5 Channels*. To examine the non-specific interaction between Halo-Ku80/Ku70 and the coverslip, control ROIs were picked that were > 6 pixels from tethered DNAs. The position of ROIs in the Cy5 channel for tethered DNA and control ROIs were identified using the calibration list and the transformation method described in *Alignment of the Cy3 and Cy5 Channels.* Stage drift was estimated using the average change in x and y position between successive frames for particles in the 532 nm channel and ROI positions were translated to compensate. In each of the 641 nm excitation frames, Cy5-labeled Halo-Ku80/Ku70 localized spots were detected using the algorithm described above and were fit to a 2D Gaussian. Integrated intensities for Cy5-Halo-Ku80/Ku70 emission signal for tethered DNA ROI was used to identify Halo-Ku80/Ku70 stoichiometry as outlined with the following method. The trajectories for Cy5 emission were first smoothed using a moving average followed by identification of the last bleaching changepoints event and median background intensity using the built-in MATLAB function ischangepts, which identifies abrupt changes in the mean that is above a specified threshold by minimization of a cost function. For each trajectory, the median background intensity was subtracted and was then normalized to the immediate state preceding complete loss of Cy5 signal. The abrupt changepoints in mean intensity were then identified in the normalized trajectory. The identified changepoints are classified as bleaching events if the following conditions are satisfied: (1) a 15 % decrease in intensity at the identified changepoint, (2) the intensity of the preceding state must be four standard deviations above the mean Cy5 background threshold. The number of bleaching events (see **Table 1**) was then used to define the number of Ku70/Ku80 molecules bound to DNA. To obtain the fractional occupancy of Ku70/80 on DNA ends plots (**Fig. 2 and Fig. S5**), for each experiment the distribution of bound Ku70/Ku80 was normalized to the number of immobilized DNA substrates. Normalized histograms (**Fig. 2B-F**) are constructed from mean fractions per occupancy bin calculated in 3 independent experiments with the standard deviation indicated by the error bars (+/-SD). Histograms of the normalized initial state intensity are constructed from initial state mean Cy5 intensity for each immobilized DNA with bound Ku70/Ku80 molecules; the initial state is defined by first identified bleaching event. Histograms of the normalized final state intensity are constructed from the immobilized DNA with a bound Ku70/Ku80 molecules using the mean final state Cy5 intensity preceding Cy5 signal loss. Histograms of the normalized bleached state intensity are constructed from the immobilized DNA with bound Ku70/Ku80 molecules using the mean intensity following Cy5 signal loss.

## Statistical analysis

Displayed uncertainties in all graph correspond to standard deviation (SD). For SIM imaging experiments, statistical analysis was performed on at least 3 independents experiments using a ratio paired t-test. For the transcription assay, a paired t-test was performed on 3 independents experiments. P-values were pictured as follow: ns p > 0.05, * p ≤ 0.05, ** p ≤ 0.01, *** p ≤ 0.001.

## Antibodies

**Table 2:** List of commercially available antibodies used in this study.

| Antibody (clone ID) | Species | Class | Reference | Manufacturer | Dilution |
| --- | --- | --- | --- | --- | --- |
| **Immunofluorescence** |  |  |  |  |  |
| Anti-Ku80 (111) | Mouse | Monoclonal | MA5-12933 | Thermo Fisher Scientific | 1:100 |
| Anti-PCNA | Rabbit | Polyclonal | Ab18197 | Abcam | 1:2000 |
| Anti-mouse 488 | Goat | Polyclonal | A11001 | Thermo Fisher Scientific | 1:1000 |
| Anti-rabbit 594 | Goat | Polyclonal | A11012 | Thermo Fisher Scientific | 1:1000 |
| Anti-mouse 647 | Donkey | Polyclonal | A-31571 | Thermo Fisher Scientific | 1:1000 |
| **Flow cytometry** |  |  |  |  |  |
| Anti-RPA32 (9H8) | Mouse | Monoclonal | Ab2175 | Abcam | 1:100 |
| Anti-PCNA | Rabbit | Polyclonal | Ab18197 | Abcam | 1:500 |
| Anti-mouse 488 | Goat | Polyclonal | A11001 | Thermo Fisher Scientific | 1:200 |
| Anti-rabbit 594 | Goat | Polyclonal | A11012 | Thermo Fisher Scientific | 1:200 |
| **Immunoblotting** |  |  |  |  |  |
| Anti-DNA-PK (18.2) | Mouse | Monoclonal | MA5-13238 | Thermo Fisher Scientific | 1:2000 |
| Anti-FBXL12 | Rabbit | Polyclonal | HPA002843 | Atlas Antibodies | 1:1000 |
| Anti-γH2AX | Rabbit | Polyclonal | 2577 | Cell Signaling Tech. | 1:1000 |
| Anti-Ku70 (N3H10) | Mouse | Monoclonal | MA5-13110 | Thermo Fisher Scientific | 1:1000 |
| Anti-Ku80 (111) | Mouse | Monoclonal | MA5-12933 | Thermo Fisher Scientific | 1:1000 |
| Anti-LIG4 | Rabbit | Polyclonal | A11432 | Abclonal | 1:1000 |
| Anti-nucleolin | Rabbit | Polyclonal | ab50279 | Abcam | 1:1000 |
| Anti-SAFA (3G6) | Mouse | Monoclonal | sc-32315 | Santa Cruz Biotech. | 1:500 |
| Anti-αTubulin | Mouse | Monoclonal | T5168 | Sigma-Aldrich | 1:25000 |
| Anti-ubiquitin | Rabbit | Monoclonal | A19686 | Abclonal | 1:1000 |
| Anti-Vinculin (7F9) | Mouse | Monoclonal | sc-73614 | Santa-Cruz Biotech. | 1:1000 |
| HRP-coupled anti-mouse | Goat | Polyclonal | 115-035-003 | Jackson ImmunoResearch | 1:10000 |
| HRP-coupled anti-rabbit | Goat | Polyclonal | 111-035-003 | Jackson ImmunoResearch | 1:10000 |
| **Immunoprecipitation** |  |  |  |  |  |
| Anti-Ku70/80 (162) | Mouse | Monoclonal | MA5-12938 | Thermo Fisher Scientific |  |
| Trueblot-HRP anti-mouse | Rat | Monoclonal | 18-8817-30 | Rockland ImmunoChemicals |  |

The anti-CtIP antibody (Clone 14-1) was a generous gift from Dr. Richard Baer. For egg extract experiments, rabbit polyclonal antibodies raised against the following *X. laevis* proteins were previously described for Ku80, XLF, XRCC4, DNA-PKcs (Graham et al., 2016), and Histone H3 (Cell Signaling). DNA-PKcs antibody was affinity purified from rabbit serum by coupling recombinant DNA-PKcs PIKK-FATC antigen (Graham et al., 2016) to AminoLink Coupling Resin (Thermo Scientific) and following manufacturer’s instructions for IgG purification. For immunoblotting, antibodies were used at the following concentrations: α-Ku80 1:10000, 1:5000 dilution of concentrated hybridoma supernatant containing α-DNA-PKcs 42-27 mouse monoclonal antibody (a kind gift of Prof. Katheryn Meek, Michigan State University), α-H3 1:500.

