## Supplementary figures with captions for "Identification of the main barriers to Ku accumulation in chromatin"

2 Equipe labélisée la Ligue contre le Cancer 2018

3 Department of Biological Chemistry and Molecular Pharmacology, Blavatnik Institute, Harvard Medical School, Boston, Massachusetts 02115

† Equal contribution.

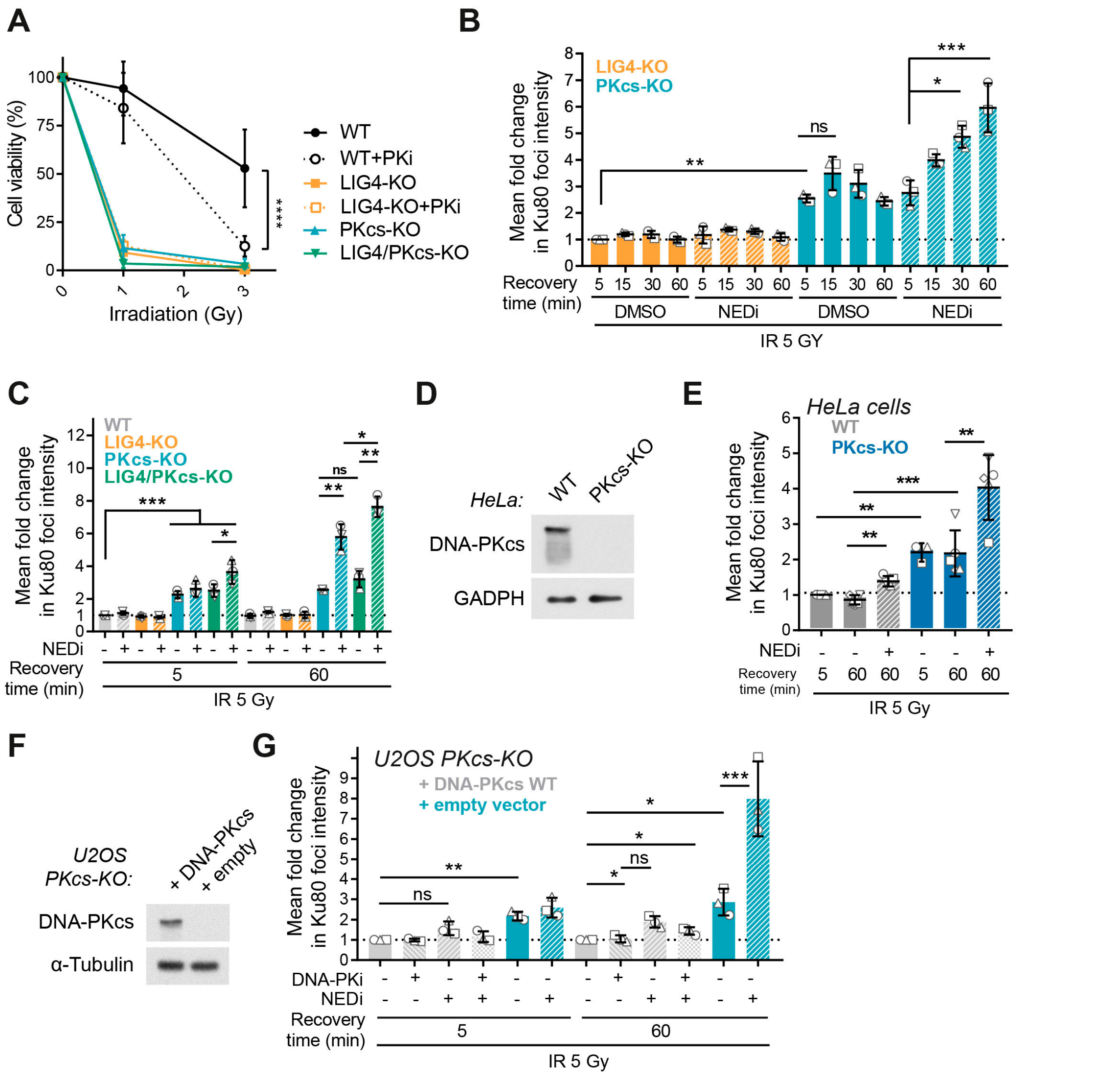


**Supplementary Figure S1: DNA-PKcs presence but not activity limits Ku accumulation at DSBs**, related to Figure 1**. A.** WT, LIG4-KO, PKcs-KO and LIG4/PKcs-KO were exposed to the indicated dose of IR, grown to 11 days, fixed and stained with crystal violet. The dye was resuspended and the absorbance of each condition measured and used as a readout of cell viability. The data corresponds to 4 independent experiments. **B.** PKcs-KO or LIG4-KO U2OS cells received 5 Gy of IR and were post-incubated for the indicated time with or without NEDi before being processed for Ku foci imaging. Ku foci average intensity was measured and normalized to the Ku foci intensity measured after 5 Gy of IR in WT U2OS to compute the fold change in Ku foci intensity in each condition, depicted on the graph. **C.** WT, PKcs-KO, LIG4-KO or LIG4/PKcs-KO U2OS cells received 5 Gy of IR and were post-incubated 5 or 60 min with or without NEDi before being processed for Ku foci imaging. Ku foci average intensity was measured and normalized to the Ku foci intensity measured after 5 Gy of IR in WT U2OS to compute the fold change in Ku foci intensity in each condition, depicted on the graph. **D.** Immunoblot of whole-cell extracts from HeLa WT or PKcs-KO. **E.** WT and PKcs-KO HeLa were treated with 5 Gy of IR and post-incubated for 5 or 60 min with or without NEDi before being processed for Ku foci imaging. Ku foci average intensity was measured and normalized to the Ku foci intensity measured after 5 Gy of IR in WT HeLa to compute the fold change in Ku foci intensity in each condition, depicted on the graph. **F.** Immunoblot of U2OS knocked-out for DNA-PKcs complemented with an empty plasmid or a plasmid expressing wild-type DNA-PKcs. **G.** U2OS PKcs-KO cells stably complemented with an empty plasmid or with a plasmid expressing WT DNA-PK were treated with 5 Gy of IR and post-incubated for 5 or 60 min with or without NEDi or PKi (3 µM NU7441) before being processed for Ku foci imaging. Ku foci average intensity was measured and normalized to the Ku foci intensity measured after 5 Gy of IR in WT U2OS to compute the fold change in Ku foci intensity in each condition, depicted on the graph.

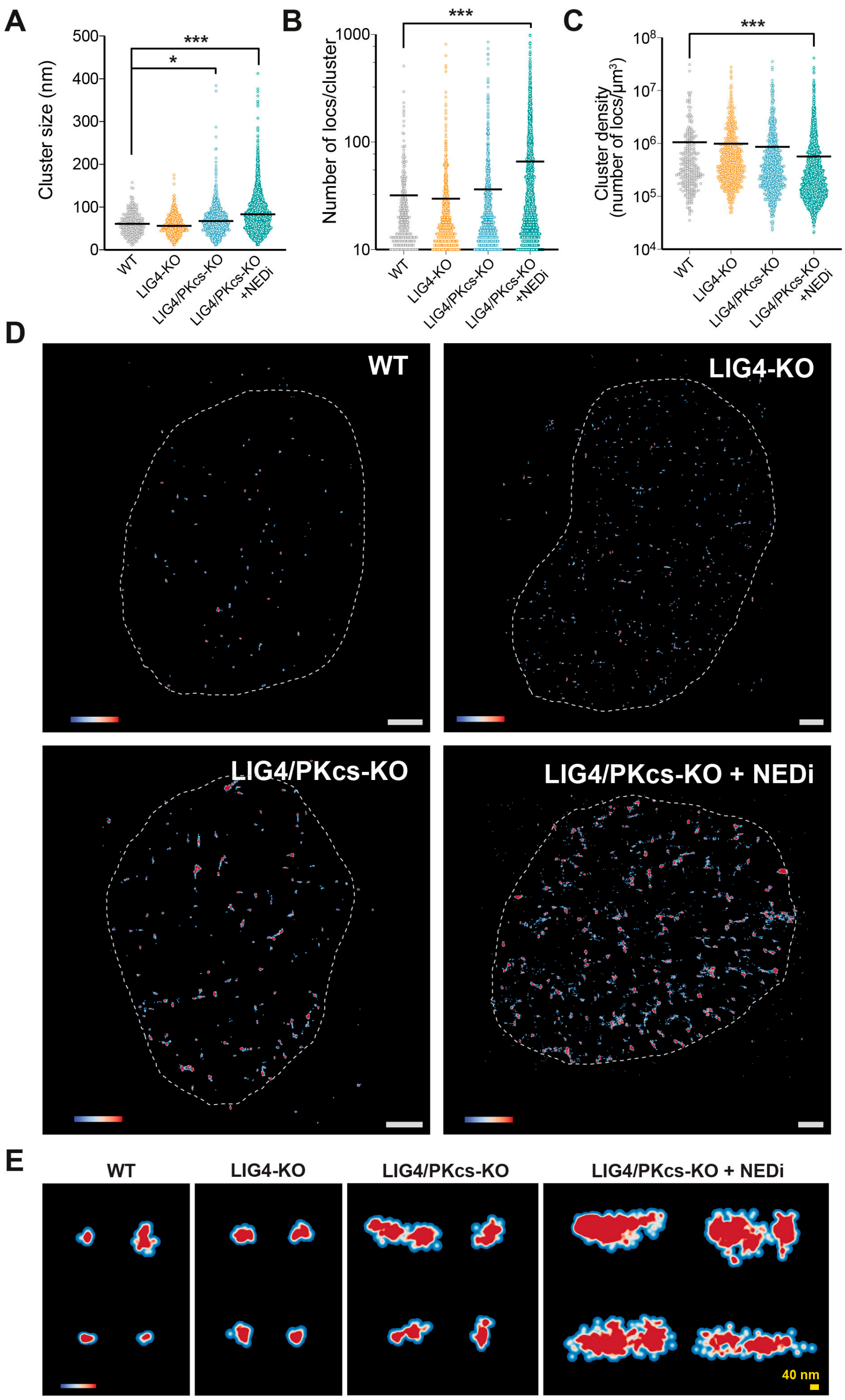


**Supplementary Figure S2: STochastic Optical Reconstruction Microscopy (STORM) of Ku foci. A-C.** Point Cloud Analyst (PoCA) analysis of STORM data was performed to estimate the size of Ku foci (A), number of localizations per cluster (B) and cluster density (C). **D**. Heat map rendering of individual localization was used to represent Ku foci in the nucleus of individual cells. Density bar, from low (blue) to high (red) localizations density. Representative images of individual nucleus are shown. Scale bar = 1 µm. **E.** Heat map rendering of individual localizations of representative clusters for each condition. Density bar, from low (blue) to high (red) localizations density. Scale bar = 40 nm.

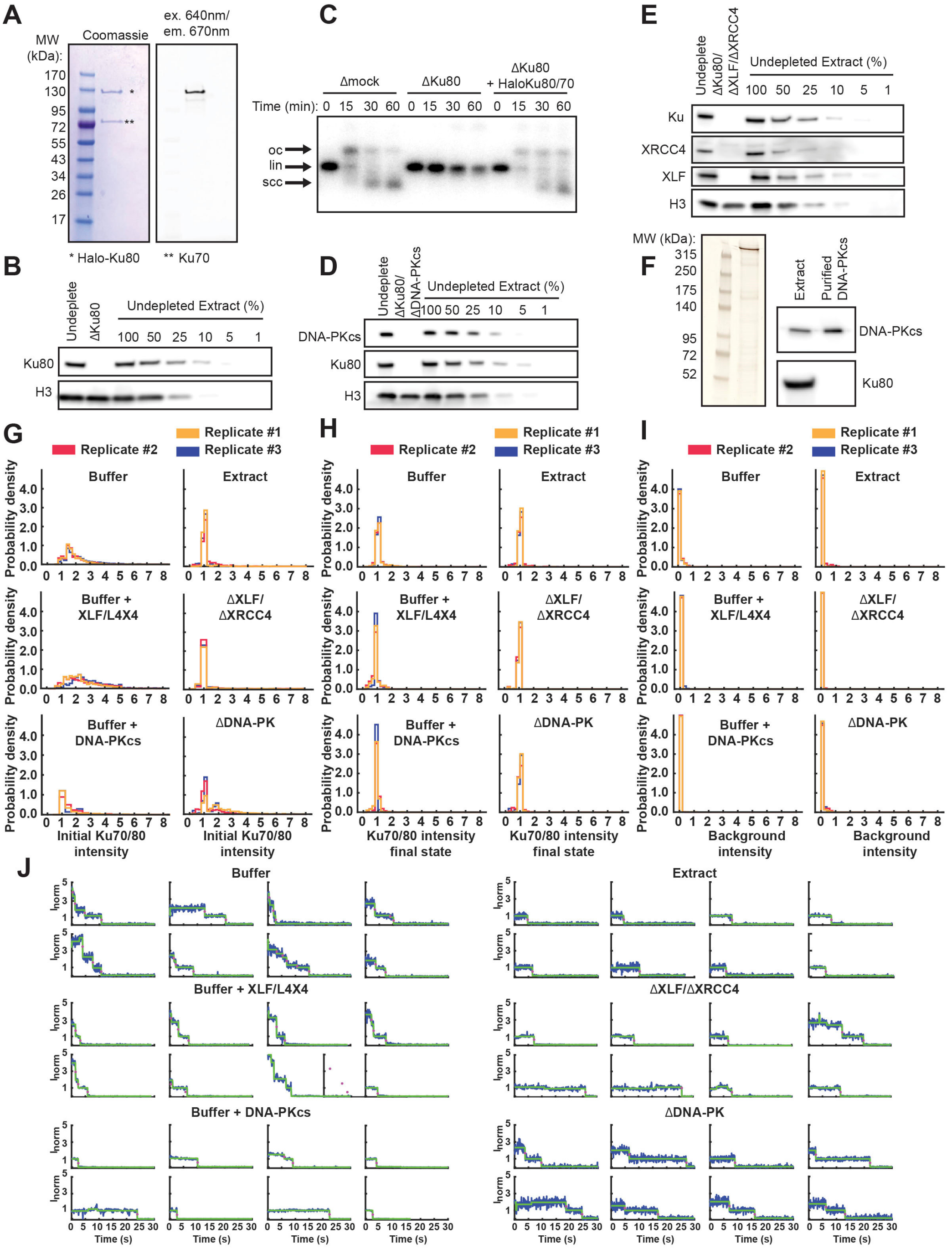


**Supplementary Figure S3: Biochemical controls, single-molecule probability distributions, and representative photobleaching trajectories** related to Figure 2**. A.** SDS-PAGE gel of purified recombinant *X. laevis* Ku80 (Halo-tagged, Cy5 labeled) and Ku70: (left) stained with colloidal Coomassie blue (InstantBlue) and (right) fluorescent emission of Cy5 label on Halo-tagged Ku80. **B.** Western blot analysis of the Ku80 immunodepleted extract. **C.** Inhibition of non-homologous end-joining by immunodepletion of Ku70/80 with αKu80 antibody and rescue with 300 nM recombinant *X. laevis* Cy5 labeled Halo-Ku80/70. Abbreviations: lin, radiolabeled linear DNA substrate; oc, open-circular products; scc, supercoiled closed-circular products. **D.** Western blot analysis of the [Ku80+DNA-PKcs] immunodepleted extracts. **E.**

Western blot analysis of the [Ku80+XLF+XRCC4] immunodepleted extracts. **F.** SDS-PAGE gel of purified endogenous *X. laevis* DNA-PKcs: (left) silver stained (Invitrogen) and (right) western blot analysis of *Xenopus* egg extract and the DNA-PKcs purified from egg extract. **G-I.** Histograms depicting the mean probability distribution for individual replicates of Cy5 Halo-Ku80/Ku70 molecules bound to 100 mer DNA for the following conditions: buffer only no extract control (Buffer), buffer plus 60 nM XLF and Lig4-XRCC4 (Buffer + XLF/L4X4), buffer plus 60 nM DNA-PKcs (Buffer + DNA-PKcs), *Xenopus* egg extract (Extract), *Xenopus* egg extract immunodepleted of XLF and XRCC4 (ΔXLF/ΔXRCC4), or *Xenopus* egg extract immunodepleted of DNA-PKcs (ΔDNA-PK). **G**. The initial intensity distribution for Ku70/80 bound to the 100 mer. **H.** The intensity distribution for final state preceding complete loss of the Cy5 signal. **I.** The background intensity distribution following complete loss of the Cy5 signal. **J.** Representative photobleaching trajectories for the following conditions: buffer only no extract control (Buffer), buffer plus 60 nM XLF and Lig4-XRCC4 (Buffer + XLF/L4X4), buffer plus 60 nM DNA-PKcs (Buffer + DNA-PKcs), *Xenopus* egg extract (Extract), *Xenopus* egg extract immunodepleted of XLF and XRCC4 (ΔXLF/ΔXRCC4), or *Xenopus* egg extract immunodepleted of DNA-PKcs (ΔDNA-PK).

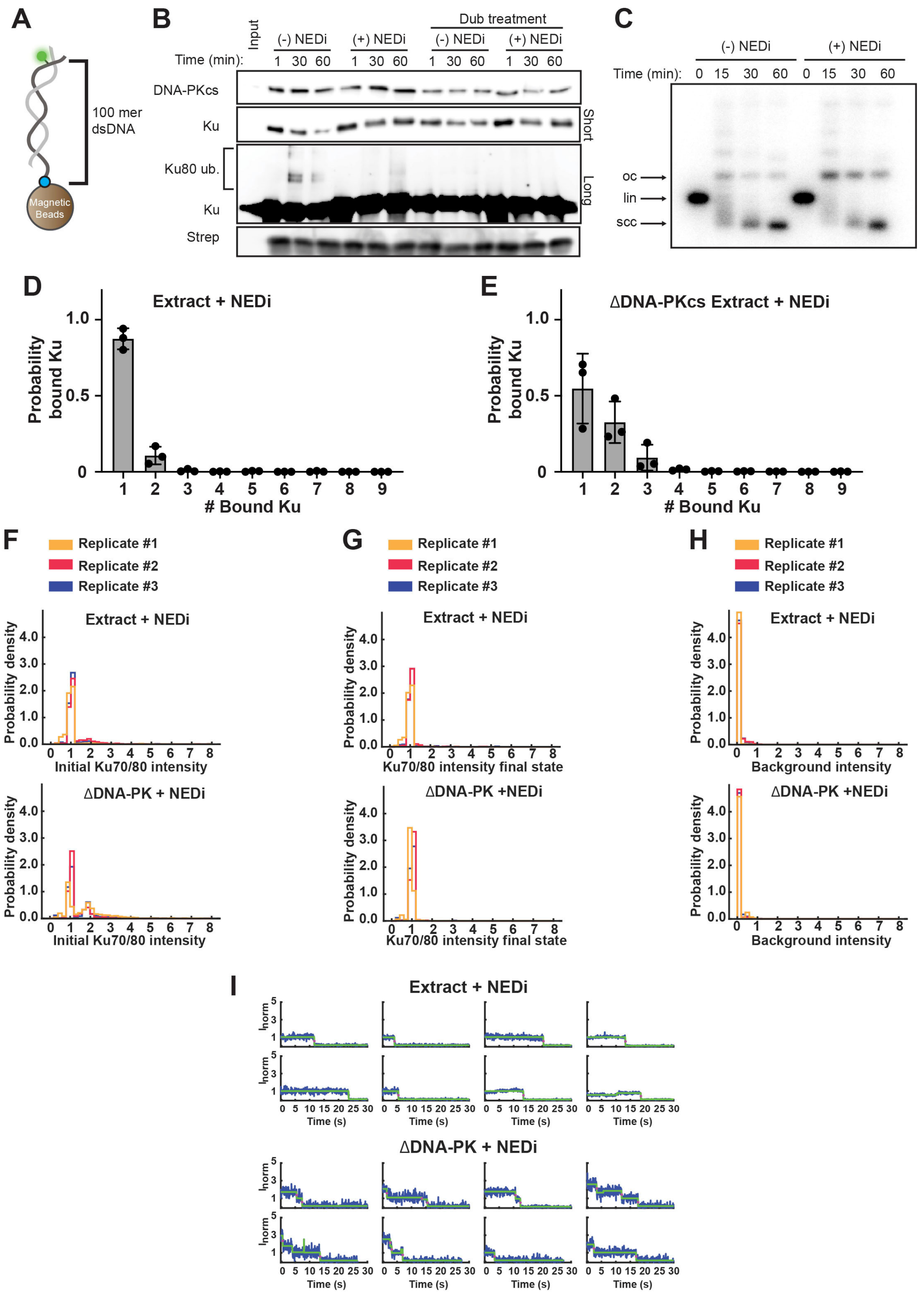


**Supplementary Figure S4: Inhibition of neddylation does not alter the Ku70/80 stoichiometry on a single DNA end in egg extract. A-B.** Inhibition of ubiquitylation of Ku80 in extract with the addition of cullin inhibitor (MLN4924; NEDi). **A.** Cartoon schematic of the DNA pulldown assay. The 5’ end of the DNA is attached to the magnetic beads. **B.** Immunoblots of NHEJ core factors (DNA-PKcs, Ku80) bound to DNA-beads over a 60 min time course, with streptavidin shown as a loading control. Input corresponds to extract diluted

1:80. Samples correspond to pulldowns of the 100 mer substrate for the following treatments: Control (DMSO only), + NEDi (200 μM neddylation inhibitor, MLN4924), as well as the same conditions de-ubiquitynated after DNA pulldown by incubation 1 h at 37°C with the purified DUB USP2. **C.** Addition of cullin inhibitor does not alter efficiency of non-homologous end-joining in egg extract. Vehicle control or neddylation inhibitor (MLN4924; NEDi) in DMSO was added to a final concentration of 200 μM. Abbreviations: lin, radiolabeled linear DNA substrate; oc, open-circular products; scc, supercoiled closed-circular products. **D-E**: On each panel the normalized histograms depicting fractional occupancy of Ku70/80 on DNA ends, constructed from mean fractions per occupancy bin calculated in 3 independent experiments. The number of events and total number of molecules observed for each experiment are reported in **Table 1**. **D.** The number of Ku molecules was monitored as described in Figure 2A**.** using *Xenopus* eggs extracts treated with 200 μM NEDi and containing Cy5-labeled Ku. **E.** The number of Ku molecules was monitored as described in Figure 2A**.** using *Xenopus* eggs extracts treated with NEDi, immunodepleted for DNA-PKcs (ΔDNA-PKcs) and containing purified Cy5-labeled Ku. **F-I.** Histograms depicting the mean probability distribution for individual replicates of Cy5 Halo-Ku80/Ku70 molecules bound to 100 mer DNA for the following conditions: *Xenopus* egg extract plus 200 μM MLN4924 cullin inhibitor (Extract + NEDi) or *Xenopus* egg extract immunodepleted of DNA-PKcs plus cullin inhibitor (Extract ΔDNA-PK + NEDi). **F**. The initial intensity distribution for Ku70/80 bound to 100 mer. **G.** The intensity distribution for final state preceding complete loss of Cy5 signal. **H.** The background intensity distribution following complete loss of Cy5 signal. **I.** Representative photobleaching trajectories for the following conditions: *Xenopus* egg extract plus 200 μM MLN4924 cullin inhibitor (Extract + NEDi) or *Xenopus* egg extract immunodepleted of DNA-PKcs plus cullin inhibitor (Extract ΔDNA-PK + NEDi).

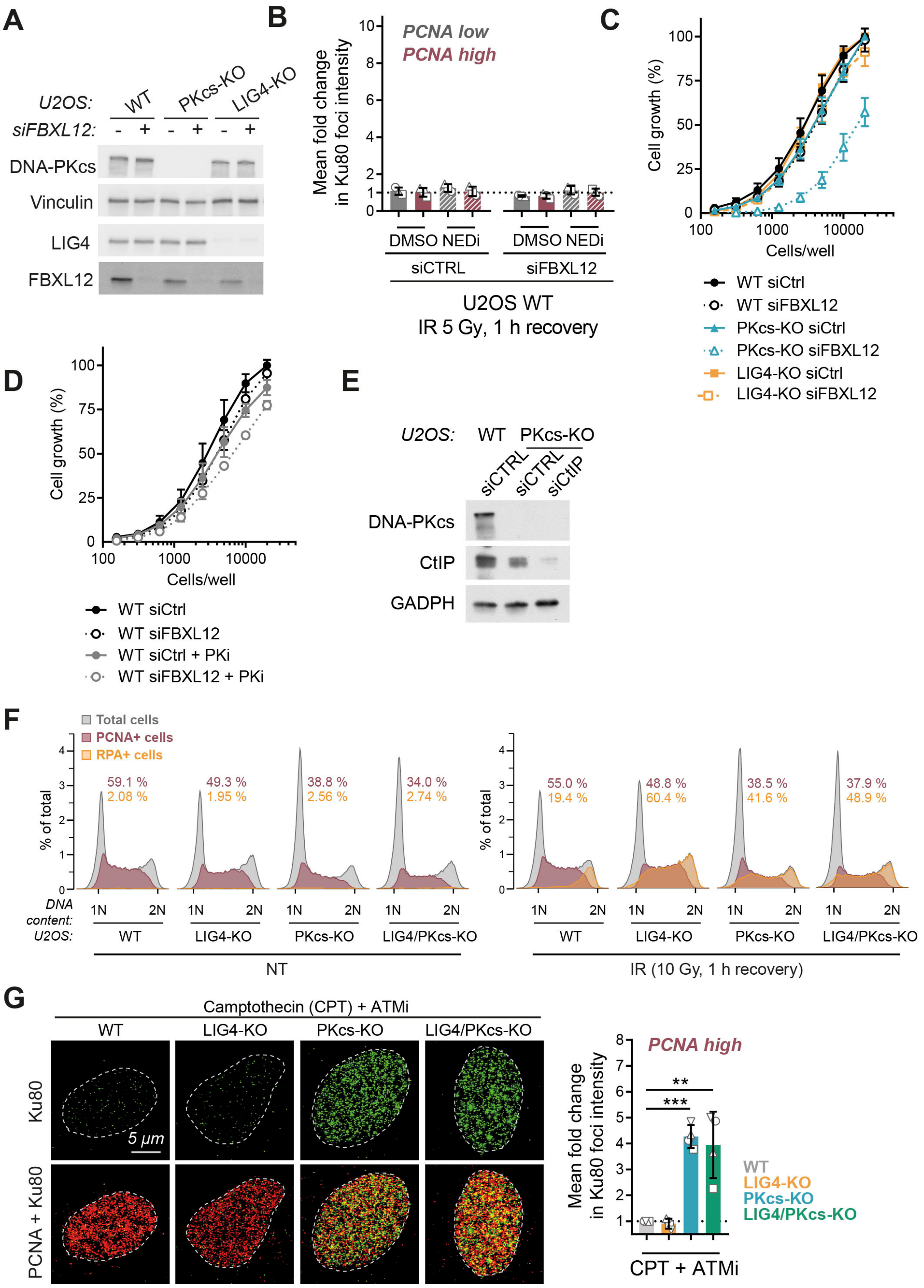


**Figure S5: Different mechanisms limit Ku loading during the cell cycle,** related to Figure 3**. A.** The depletion of FBXL12 in U2OS WT, PKcs-KO or LIG4-KO transfected by control (CTRL) or anti-FBXL12 siRNA was confirmed by immunoblotting in whole-cell extracts. **B.** U2OS WT which had been transfected by siRNA control or against FBXL12 were pre-treated or not with NEDi, received 5 Gy of IR and were post-incubated 1 h before being processed for immunofluorescence. A PCNA staining was used to identify the cells in S-phase. Ku foci average intensity was measured in each condition and normalized to the Ku foci intensity in U2OS WT measured 5 min after 5 Gy of IR to compute the fold change in Ku foci intensity in each condition, displayed on the graph. **C.** Cell fitness was evaluated by monitoring the proliferation of U2OS WT, PKcs-KO or LIG4-KO after transfection with Ctrl or anti-FBXL12 siRNAs. Three days after plating the amount of cells in each well was evaluated using sulforhodamine B (SRB) staining. **D.** Cell fitness was evaluated by monitoring the proliferation of U2OS WT after transfection with Ctrl or anti-FBXL12 siRNAs. Three days after plating with or without 0.5 µM DNA-PK inhibitor nedisertib the amount of cells in each well was evaluated using SRB staining. **E.** Flow cytometry analysis of RPA32 signal (yellow) in WT, PKcs-KO, LIG4-KO or LIG4/PKcs-KO U2OS cells irradiated with 10 Gy. PCNA signal (red) was measured to identify cells in S-phase. **F.** The depletion of CtIP in U2OS WT or PKcs-KO transfected by control (CTRL) or anti-CtIP siRNA was confirmed by immunoblotting in whole-cell extracts. **G**. WT, PKcs-KO, LIG4-KO or LIG4/PKcs-KO U2OS cells were pre-incubated 1 h with ATMi, then treated with 1 µM CPT for 1 h before being processed for Ku foci imaging. A PCNA staining was used to identify the cells in S-phase. Representative pictures are shown on the left panel. Ku foci average intensity was measured and normalized to the Ku foci intensity measured in WT U2OS to compute the fold change in Ku foci intensity in each condition, depicted on the right panel.
